# An Oral Combination therapy against SARS-CoV-2 based on Synergistic Action of Auranofin and Remdesivir

**DOI:** 10.64898/2025.12.19.695342

**Authors:** Oyahida Khatun, Rohan Narayan, Santhosh Kambaiah Nagaraj, Rishad Shiraz, Abinaya Kaliappan, Sumandeep Kaur, Ankur Singh, Shridhar Narayanan, Radha Krishan Shandil, Shailly Tomar, Shashank Tripathi

**Author notes:** These authors contributed equally (Oyahida Khatun, Rohan Narayan, Santhosh Kambaiah Nagaraj).

## Abstract

The combination of direct-acting and host-directed antivirals targeting SARS-CoV-2 represents an attractive treatment strategy to combat COVID-19. In our previous work, we showed that the FDA-approved anti-arthritis drug Auranofin restricts SARS-CoV-2 replication and pathology in an animal model. Here, we report that Auranofin inhibits SARS-CoV-2 by targeting viral entry and main protease (Mpro) activity without affecting viral transcription. Time-of-addition studies combined with functional assays of viral entry, protease activity, and cell–cell fusion delineated its inhibitory effects at both early and late stages of the viral life cycle. Molecular docking and isothermal titration calorimetry analyses of Auranofin and the known Mpro inhibitor Nirmatrelvir indicated competitive binding within the Mpro active-site pocket. In addition, Auranofin attenuated NF-κB-dependent signalling and suppressed proinflammatory cytokine production. Combination studies with SARS-CoV-2 targeting nucleoside analogues, revealed the strongest synergistic antiviral activity with remdesivir in vitro. Comparable synergy was observed between auranofin and GS-621763, the orally bioavailable derivative of remdesivir, and was further validated in a preclinical animal model. Collectively, these findings provide a rationale for the further development of auranofin-nucleoside analog combinations targeting SARS-CoV-2.

**Funding:** This research has been supported by ICMR (IIRPIG-2023-0000978) and BIRAC grant (BT/CS0070/06/22) to ST. We acknowledge the infrastructure and research support provided to IISc by the Crypto Relief Fund, L&T Trust, DST-FIST program, Institute of Eminence Fund, Ministry of Education, and the DBT-IISc partnership program (Phase II). RN acknowledges DBT-RA fellowship, SK acknowledges PMRF fellowship, and RS acknowledges FICCI-PMRF fellowship.

**Research in context:** *Evidence before this study:* Multiple previous studies have provided evidence showing that auranofin is an antiviral agent targeting SARS-CoV-2 through a host-targeting mechanism, involving the inhibition of thioredoxin reductase and redox homeostasis. Mixed results have been reported regarding the effect of this drug on inhibiting virus-induced syncytia. A prior study from our lab demonstrated the drug’s effectiveness in reducing SARS-CoV-2 viral loads in both cell and animal models.

*Added value of the study:* Our study firmly establishes the synergistic use of Auranofin and Remdesivir GS 621763 in the treatment of SARS-CoV-2 infection.

## Introduction

COVID-19 is a respiratory illness caused by Severe Acute Respiratory Syndrome Coronavirus-2 (SARS-CoV-2), an enveloped single-stranded positive-sense RNA virus.^1^ Since the initial cases were reported in December 2019, the COVID-19 pandemic has resulted in the deaths of over 7.1 million people globally, as of November, 2024.^2^ Mortality rates were much higher among older adults and those with underlying health conditions, and much of the pathobiology of COVID-19 was due to a combination of direct viral injury to tissues and aberrant immune responses against infection.^3,4^ Fortuitously, the introduction of a plethora of vaccines and effective antivirals had a significant impact on reducing the global morbidity and mortality associated with COVID-19^5^. Antivirals play a very important role in complementing vaccine efficacy by reducing viral load and thereby mitigating virus transmission.^6^ Direct-acting antivirals (DAA) against SARS-CoV-2 may target virus entry, virus-encoded proteases, or viral genome replication and transcription. These classes of drugs generally pose the risk of giving rise to the emergence of drug-resistant mutants.^7,8^ In contrast, Host-Directed Antivirals (HDA) target host components that are crucial during the virus lifecycle, and in the context of SARS-CoV-2, include the immunosuppressive humanised monoclonal antibody drug Tocilizumab ^9^, anti-inflammatory drugs and immunomodulators. Synergistic antiviral effects against viral diseases can be achieved through combination treatment with different classes of drugs that have independent mechanisms of action and act upon different stages of the virus lifecycle.^11^ Such synergistic intervention strategies have been reported previously for a few viruses including SARS-CoV-2.^12–14^

The SARS-CoV-2 Mpro is a cysteine protease responsible for most cleavage events that occur in the virus’s precursor polyprotein. Mpro achieves protein cleavage via the catalytic dyad His41 and Cys145. The key active site residues critical for substrate binding and catalysis include Cys145, His41, Glu166, His163 and Gly143.^15,16^ The structure and function of Mpro is conserved across Coronaviruses and there is no human homolog of this protein, thus making it an ideal antiviral target.^17^ Nirmatrelvir was the first protease inhibitor specifically developed to target SARS-CoV-2 Mpro and is administered to patients in combination with Ritonavir for the treatment of COVID-19.^18,19^ Our previous work showed that the FDA-approved drug Auranofin, used for the treatment of rheumatoid arthritis, shows antiviral activity against SARS-CoV-2 in both cell culture and animal models.^20^ Here, we provide evidence that auranofin affects virus entry and displays synergistic antiviral activity with remdesivir and GS-621763, an orally bioavailable prodrug of the remdesivir parent nucleoside GS-441524, supporting combination therapeutic strategies targeting SARS-CoV-2.

## Results

### Auranofin inhibits SARS-CoV-2 entry and Mpro activity

To understand the mode of action of Auranofin against SARS-CoV-2, we first tested its effect on virus entry using luciferase-expressing non-replicating SARS-CoV-2 spike pseudotyped virus particles.^21^ For this, HEK293T cells expressing human ACE2 receptor (HEK293T-ACE2) cells were pre-treated with 1 µM Auranofin for 3 h, followed by spike pseudotyped virus infection in the presence of the drug for 48 h. The Luciferase activity was measured 48 h post-infection (48 hpi) (Fig 1A). We observed that Auranofin treatment reduced virus entry compared to virus control, as shown by a 3-fold decrease in luciferase signal (Figure 1B). To validate this finding, we performed a time-of-addition (ToA) assay where HEK293T ACE2 were pre-treated with 1 µM Auranofin for 3 h and infected with 10 MOI SARS-CoV-2 (Hong Kong/VM20001061/2020 isolate) (hereby referred as SARS-CoV-2) in the presence of the drug (Figure 1 C). After 3 h of infection, cell lysates were collected and analysed by Western blot. Auranofin treatment significantly reduced virus SARS-CoV-2 spike protein expression, indicating inhibition of virus entry (Figure 1 D, E).

**Figure 1.**
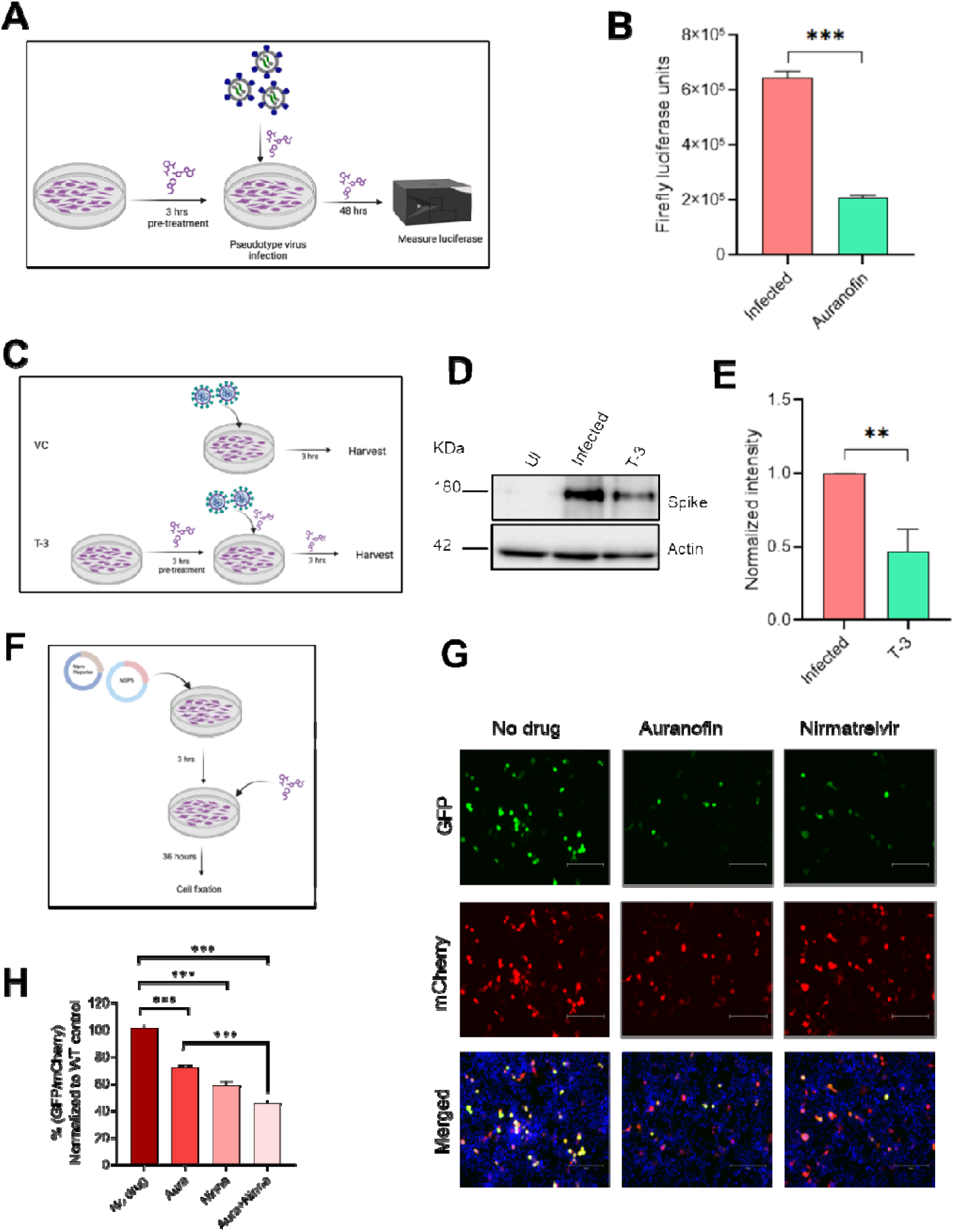
Auranofin inhibits SARS-CoV-2 entry and viral Mpro activity in cell culture. (A) Schematic of pseudotype virus entry. HEK293T ACE2 cells pretreated with 1 μM Auranofin were infected with SARS-CoV-2 spike pseudotyped particles and 3 hours post-infection (hpi) luciferase activity was measured. (B) Graph showing firefly luciferase activity for infection control and Auranofin-treated conditions. (C) Schematic of time-of-addition (ToA) assay. HEK293T ACE2 cells pretreated for 3 h with 1 μM Auranofin were infected with 10 MOI SARS-CoV-2 in the presence of drug (T-3). After 3 h, cells were harvested for western blot analysis. (D) Western blot showing expression of SARS-CoV-2 spike upon Auranofin treatment, compared to infected control (VC) and (E) quantification of band intensity. (F) Schematic of SARS-CoV-2 FlipGFP^Mpro^ assay. HEK293T cells were co-transfected with FlipGFP^Mpro^ reporter construct and SARS-CoV-2 NSP5 WT plasmid, 1 μM Auranofin was added after 3 h and 36 h later cells were fixed and GFP expression was analyzed under a fluorescence microscope. Treatment with Nirmatrelvir served as a positive control. (G) Representative fluorescence images showing differences in GFP fluorescence intensity in 3 different conditions as indicated. (H) Total GFP and mCherry signals from each field were quantified using ImageJ/Fiji. The GFP signals were normalized to the mCherry signal, and then the GFP values for each treatment condition were normalized to the no-drug control. All data are from 3 independent biological replicates. ∗∗p < 0.01, ∗∗∗p < 0.001; ns, non-significant using two-tailed unpaired t-test or Welch ANOVA with Dunnett’s T3 multiple comparison tests, where applicable. Error bars represent mean ± SD.

After viral entry, the next step is the translation of the incoming viral genome into polyprotein, followed by processing by viral proteases Mpro (or 3CLpro; also called the main protease) and PL^pro^ (papain-like proteases).^22,23^ To understand the effect of Auranofin on SARS-CoV-2 main protease activity, we used a previously reported FlipGFP^Mpro^ assay.^24^ Briefly, the assay leverages a Green Fluorescent Protein (GFP) based reporter system to detect the activity of the viral protease. The reporter assay is based on the principle that the SARS-CoV-2 Mpro, encoded by the virus’s NSP5, can cleave a substrate, leading to the reassembly of the split-GFP and thereby increasing fluorescence.^24^ To optimise the reporter system, HEK293T cells expressing the FlipGFP^Mpro^ reporter construct were transfected with plasmids encoding the SARS-CoV-2 NSP5 wild type (WT) or the catalytically inactive mutant NSP5 C145A. Analysis of fluorescence activity 24 h post-transfection revealed higher GFP intensity in cells expressing the wild-type NSP5 construct compared to the weak signal observed in cells expressing the NSP5 mutant, demonstrating the optimal efficiency of the reporter assay (Sup Fig 1). To analyse the effect of Auranofin on SARS-CoV-2 Mpro activity, HEK293T cells were co-transfected with FlipGFPMpro reporter, and NSP5 WT plasmids and 3 h later, the media were replaced with fresh complete DMEM containing 1 µM Auranofin or 7.8 μM Nirmatrelvir, a known SARS-CoV-2 Mpro inhibitor used here as a positive control. After 36 h, cells were fixed, and GFP fluorescence was analysed using a fluorescence microscope (Fig. 1F). We observed a ∼ 40% reduction in GFP intensity in the presence of Auranofin suggesting that Auranofin significantly inhibits the SARS-CoV-2 Mpro activity (Fig 1 G, H).

To further delineate the effects of auranofin on SARS-CoV-2 entry, we performed virus entry assays in HEK293T ACE2 cells and analyzed drug-treated cells by confocal microscopy. Auranofin treatment resulted in a significant reduction in colocalization between SARS-CoV-2 spike protein and EEA1-labelled early endosomes (Fig. 2A, B), indicating impaired trafficking of incoming viral particles to early endosomal compartments. Consistent with this, spike-associated fluorescence intensity was significantly reduced upon treatment with 1 µM auranofin (Fig. 2C), suggesting decreased viral internalisation and/or altered post-entry trafficking. Uninfected cells treated with Auranofin showed a dispersed phenotype of EEA-1 positive endosomes across the cytoplasm, compared to a predominantly perinuclear localisation in control (Sup Fig 1). Together, these data support a role for auranofin in modulating endosomal entry or early endosomal trafficking of SARS-CoV-2 particles. The observed effects on virus entry were not due to impaired attachment of virus onto the cell surface, as demonstrated by results from the virus binding assay, which showed no reduction of cell surface staining of spike in auranofin-treated conditions (Fig. 2D, E).

**Figure 2.**
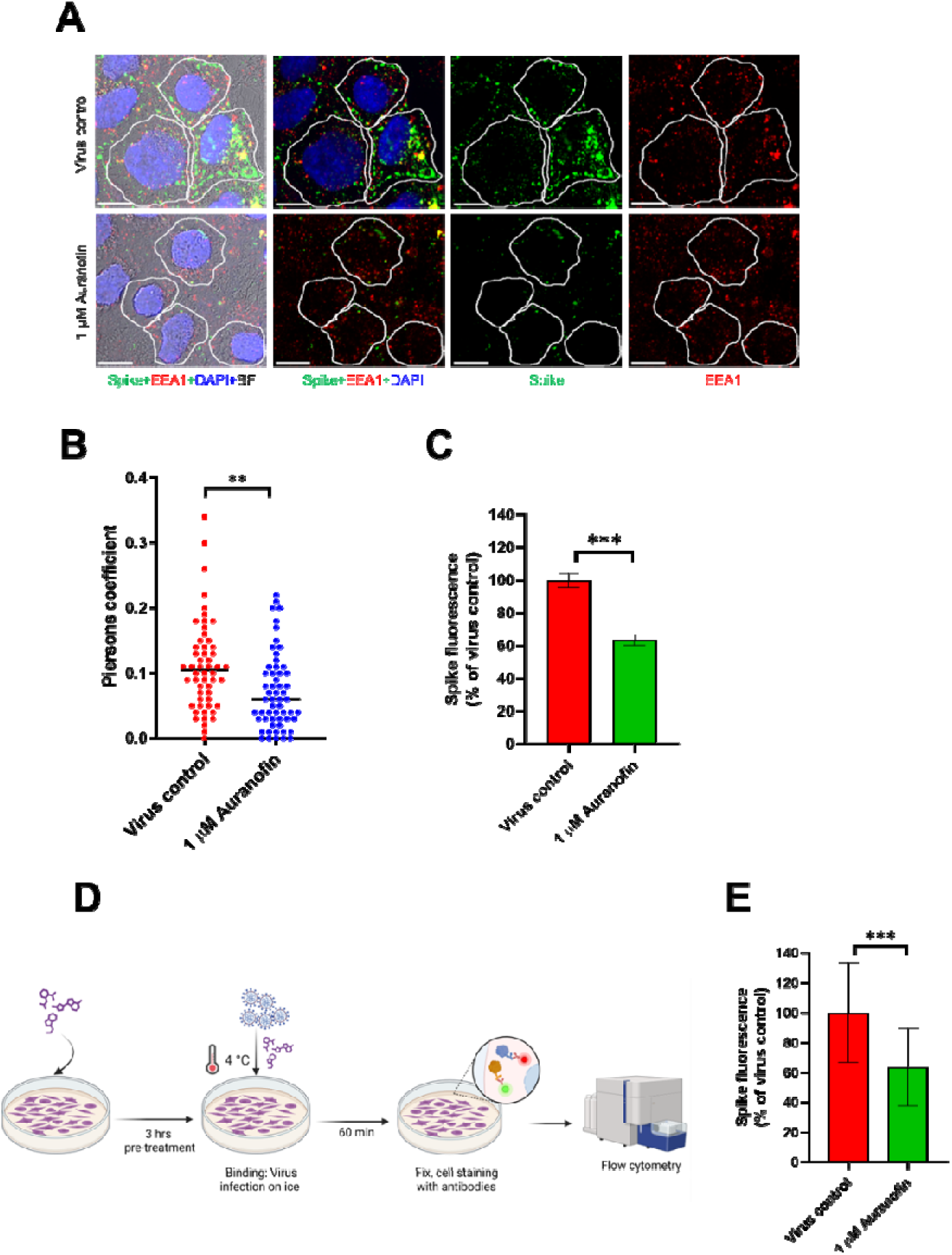
Auranofin impairs endosomal entry of SARS-CoV-2 without affecting viral attachment. (A-C) HEK293T ACE2 cells pre-treated with 1 µM auranofin were infected with 10 MOI SARS-CoV-2 on ice for 1 h and moved to 37°C in the presence of drug for 20 min. Cells were then fixed and labelled with antibodies against spike and EEA-1. (A) Confocal microscopy images with virus spike and EEA-1 positive early endosomes are shown in green and res respectively. Individual cells are indicated by ROIs drawn along with cell boundaries using bright field images as reference. (B) Colocalization between spike and EEA-1 positive vesicles was quantified using Imagej/Fiji, and piersons coefficients are plotted from an average of 10 ROIs per image field. Fluorescence intensity of spike positive vesicles quantified from green channel is shown in (C). (D-E) Schematic for protocol used in virus binding assay is shown in (D). HEK293T ACE2 cells were pre-treated with 1 µM auranofin and infected with 50 MOI SARS-CoV-2 on ice for 1 h. Cells were then washed, paraformaldehyde fixed and labelled with antibodies against virus spike. Total number for spike positive cells was quantified by flow cytometry, shown in (E). (A-C) are from 3 biological replicates and (D-E) are from 2 repeats. ∗p < 0.5, ∗∗p < 0.01, ∗∗∗p < 0.001; ns, using two-tailed unpaired t-test or Welch ANOVA with Dunnett’s T3 multiple comparison tests, where applicable. Error bars in represent mean ± SD.

### Effect of Auranofin on intermediate and late stages of virus infection

After incoming viral genome translation and polyprotein processing, the viral polymerase assembles and starts transcription and replication of the viral genome.^25^ We first investigated the possible effects of auranofin on the SARS-CoV-2 RNA-dependent RNA polymerase (RdRp) activity using a previously reported replicon-based dual-luciferase reporter assay.^26^ The systems involved co-transfecting cells with a bicistronic reporter construct p(+)RLuc–(−)UTR–FLuc and a plasmid expressing the SARS-CoV-2 RdRp. The bicistronic reporter expresses Renilla luciferase as a host-driven internal control, while firefly luciferase is expressed only following the virus RdRp-dependent conversion of a negative-sense viral RNA into a translatable positive-sense RNA.^26^ Experimentally, the drugs auranofin or remdesivir were added to cells 6 h post-transfection and luciferase activity was analysed after 48 h. No effect of auranofin on virus RdRp activity was observed, compared to remdesivir which served as positive control (Fig 3A). Next, we explored the effects of Auranofin on viral transcription by performing a time-of-addition assay (Fig 3 B). We observed that treatment of HEK293T ACE2 cells with Auranofin 6 hpi did not inhibit virus infection as shown by similar SARS-CoV-2 spike protein expression in control and treated conditions (Fig 3 C,D). No effect was observed in vRNA levels, as shown by qRT-PCR-based quantification (Fig. 3E), indicating that Auranofin does not inhibit virus polymerase activity and late translation.

**Fig 3.**
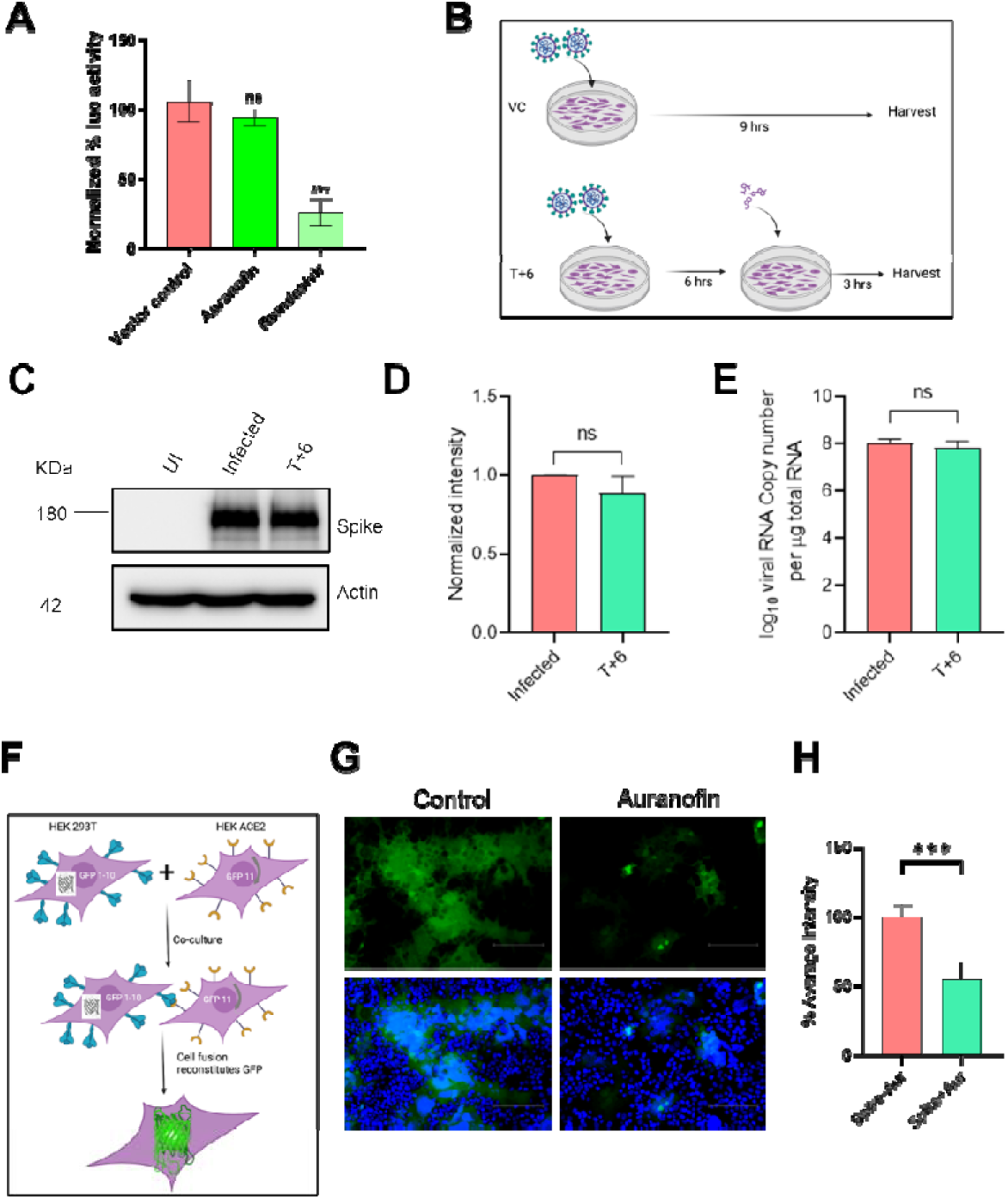
Auranofin does not inhibit SARS-CoV-2 virus polymerase activity but reduces virus-induced syncytia formation. (A) Schematic of time-of-addition assay. HEK293T ACE2 cells were infected with 10 MOI SARS-CoV-2 and 6 hpi treated with 1 μM Auranofin (T+6). Cells were harvested after a further incubation of 3 h. (B) Western blot expression of SARS-CoV-2 spike upon Auranofin treatment, compared to infected control, and (C) quantification of band intensity. (D) SARS-CoV-2 viral RNA load from infected cells, quantified by qRT PCR. (E) Schematic showing the split-GFP-based reporter system for studying cell-cell fusion assay. HEK293T cells expressing SARS-CoV-2 spike and GFP 1-10 were pre-treated with 1 µM Auranofin and co-cultured with HEK293T ACE2 cells expressing GFP 11. After 24 h, cells were fixed, nuclei labeled with DAPI, and images were captured using a fluorescence microscope. (F) Shows representative fluorescence microscopy images and (G) shows average GFP intensity from different fields quantified using a CX4 High Content Screener. (E) is from 1 experiment. All other data are from 3 independent biological replicates. ∗∗∗p < 0.001; ns, non-significant using two-tailed unpaired t-test. Error bars represent mean ± SD.

One of the very late events of SARS-CoV-2 infection in permissive cells is syncytia formation, which is characterised by cell-cell fusion mediated by the virus spike protein, resulting in the formation of multinucleated giant cells.^27,28^ To test the effects of Auranofin on virus-induced cell-cell fusion, we used a split-GFP complementation system, where GFP 1-10 and GFP 11 are expressed in two separate cell populations, generating fluorescence signals only upon fusion.^28^ HEK293T cells expressing SARS-CoV-2 spike and GFP 1-10 pre-treated with 1 µM Auranofin were co-cultured with HEK293T-ACE2 cells expressing GFP 10-11 in the presence of the drug. Analysis of GFP fluorescence after 24 h showed ∼50% reduction of SARS-CoV-2-induced syncytia formation in the presence of Auranofin (Fig 3 F-H). Taken together, the data demonstrate that Auranofin does not interfere with post-entry stages like SARS-CoV-2 replication and transcription or translation. In contrast, Auranofin can impair late cell–cell fusion events associated with virus spread. SARS-CoV-2 syncytia formation aids in virus cell-cell transmission and, therefore, inhibition of this process is a potential antiviral target. ^29,30^

### Effect of Auranofin on NFκB pathway

Auranofin is an FDA-approved anti-inflammatory drug used in the treatment of rheumatoid arthritis.^31^ We have previously reported the antiviral efficacy of Auranofin in mitigating SARS-CoV-2 infection in Syrian golden hamsters, concomitantly suppressing proinflammatory IL-6 production in the lungs.^20^ Hence, we examined the effects of this drug on the NFkB inflammatory pathway. Towards this, we used a well-established dual luciferase assay, which used HEK293T cells co-transfected with a plasmid expressing firefly luciferase driven by NFκB promoter, a constitutively active Renilla luciferase-expressing plasmid pRLTK, and an empty vector. After 24 hours, these cells were treated with Auranofin and induced with TNF-alpha to activate the NF-kB pathway in the presence of the drug (Sup Fig 2A). Luciferase activity analyzed after 12 h revealed ∼30% inhibition of the NFkB pathway upon treatment with Auranofin (Sup Fig 2B). Further on, we also tested the effects of Auranofin in the context of SARS-CoV-2 infection and observed over 3-fold decrease in NFkB activation in the presence of Auranofin (Sup Fig 2C). This was also supported by a ∼2-fold reduction in TNF alpha expression (Sup Fig 2D). These data confirmed the anti-inflammatory effect of Auranofin during SARS-CoV-2 infection.

### Auranofin directly binds to SARS-CoV-2 Mpro to inhibit its activity

The data presented so far indicate that Auranofin treatment affects the SARS-CoV-2 life cycle by abrogating virus Mpro activity, in addition to inhibiting virus entry, syncytia and the host NFkB inflammatory pathway. To better understand the direct effects of Auranofin on SARS-CoV-2 Mpro, we examined the in-silico binding of both Auranofin and Nirmatrelvir to SARS-CoV-2 Mpro by performing guided structural docking analysis. Both compounds showed stable docking with Mpro near the protein catalytic site; however, Nirmatrelvir demonstrated stronger binding to Mpro, with a binding free energy of −9.72 kcal/mol, which was ∼1.4-fold lower than that of Auranofin (−7.13 kcal/mol) (Fig a A-C). Structural analysis of the docked complexes highlighted critical hydrogen-bonding interactions between Auranofin and the key catalytic dyad residues (His41 and Cys145) of Mpro (Fig 4 A,B Sup Fig 3 A-B). Next, we proceeded to validate these findings using Isothermal Titration Calorimetry (ITC) which directly measures the heat changes associated with ligand-protein binding interactions. The dissociation constants (K_D_) for the Mpro-Auranofin and Mpro-Nirmatrelvir interactions were determined to be 197 ± 16.6 μM and 25 ± 2.5 μM, respectively (Fig 4 D,E). In contrast, the KD for the interaction between the Mpro active site mutant (C145A) and Auranofin was measured at 555 ± 127 μM, approximately a three-fold increase compared to the native Mpro (Fig 4 F). The purity and molecular weight of Mpro and the active site mutant was confirmed by SDS-PAGE (Sup Fig 3C). These ITC findings suggest that both compounds primarily bind within the active site pocket of Mpro.

**Fig 4.**
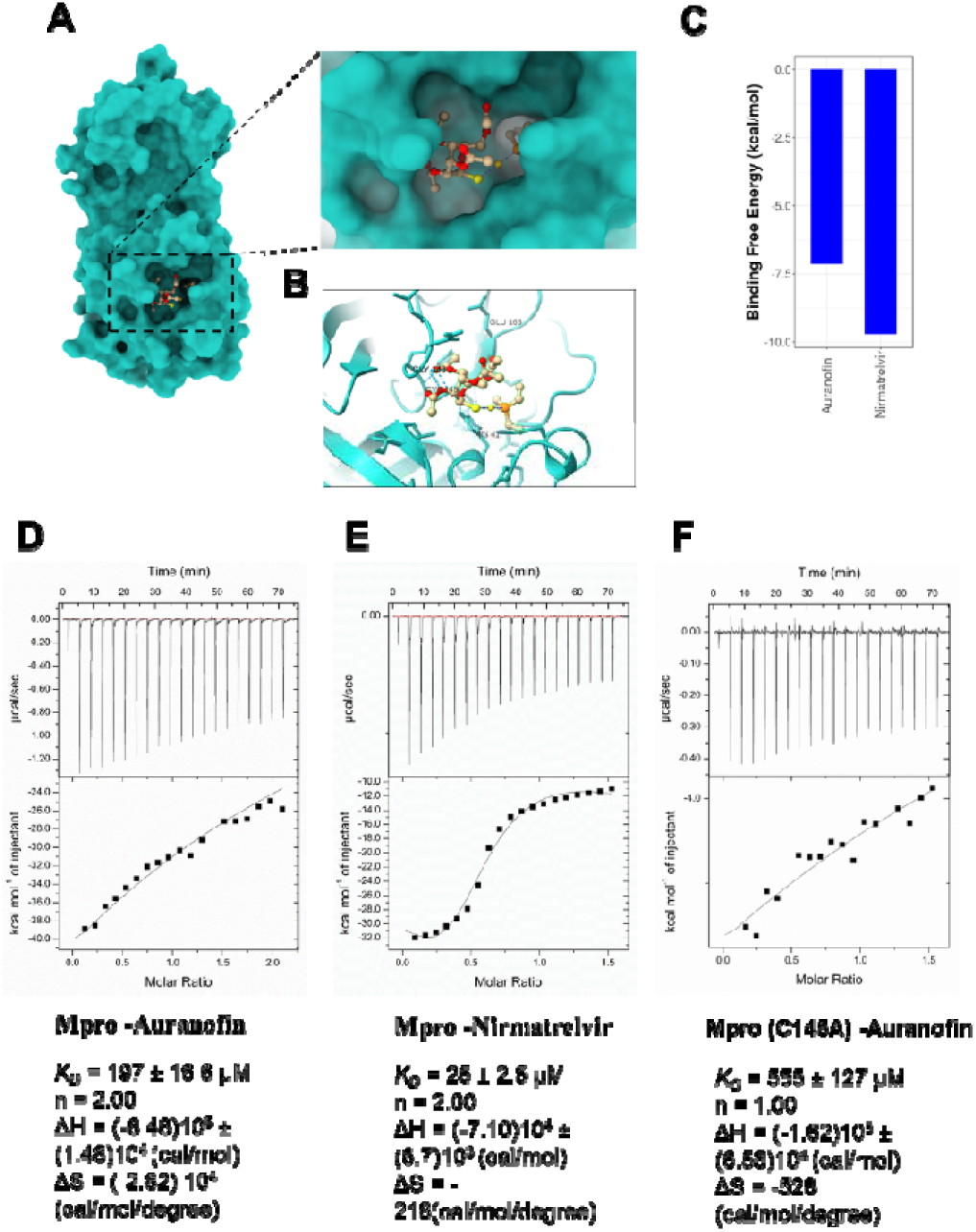
Auranofin directly binds to SARS-CoV-2 Mpro to inhibit its activity. Binding free energy and docking of Auranofin against SARS-CoV-2 Mpro were performed using Autodock. (A-B) Docked structures of SARS-CoV-2 Mpro to Auranofin and corresponding Hydrogen bonding interactions with key amino acids are shown in (A) and (B) respectively. (C) Shows binding free energy for Auranofin and Nirmatrelvir. (D-F) ITC Sensogram and isothermodynamics for binding interactions between Mpro and auranofin and nirmatrelvir are shown in (D) and (E) respectively. (F) shows similar data set for interactions between the catalytic mutant of Mpro C145A and auranofin.

When Nirmatrelvir was titrated against the preincubated Mpro-Auranofin complex, its affinity for Mpro decreased nearly four-fold, resulting in a KD of ∼102 ± 4.8 μM (Sup Fig 3 D). Conversely, Auranofin exhibited minimal binding (KD = ∼909 ± 34 μM) when titrated against the preincubated Mpro-Nirmatrelvir complex (Sup Fig 3E), indicating that Nirmatrelvir partially outcompetes Auranofin for Mpro binding. The ΔH values derived from the ITC isotherms indicate that the binding interactions are exothermically driven (Sup Fig 3 D,E). We also tested the binding of Auranofin to purified SARS-CoV-2 Spike RBD but detected no interaction, with a KD of 1135 ± 84 µM (Sup Fig 4 A). The purity and molecular weight of spike RBD was confirmed by western blot (Sup Fig 4 B).

Additionally, we assessed whether auranofin affects spike structural integrity or oligomerization by incubating the drug with cells which were either transfected to express SARS-CoV-2 spike protein or infected with live virus to express the protein. Non-reducing PAGE analysis of virus spike in both cases saw the protein migrate as a single species corresponding to its expected molecular weight, with no additional bands or mobility shifts detected, indicating that auranofin does not induce gross alterations in Spike oligomeric state or native conformation following de novo synthesis or during infection (Sup Fig 4C,D). In summary, the ITC titration curves demonstrate that the identified ligands bind spontaneously to Mpro with favorable thermodynamic parameters, while sequential binding experiments highlight their ability to mutually inhibit binding affinity at their respective binding sites.

### Auranofin exhibits synergistic antiviral effects with nucleoside analogs in cell culture model

After establishing the mode of action of Auranofin which is a combination of effects on early steps of SARS-CoV-2 infection as well as inhibition of inflammatory immune signaling, next we examined the potential synergistic or additive effect of this drug with DAAs. To this end, we tested Auranofin in combination with widely established SARS-CoV-2 DAAs namely nucleoside analogues Remdesivir ^32^, Molnupiravir ^33^, and the SARS-CoV-2 Mpro inhibitor Nirmatrelvir.^18^ We first calculated the IC90 of the drugs, including Auranofin using SARS-CoV-2 recombinant infectious clone with Nanoluciferase gene hereby referred to as SARS-CoV-2 Luc^34^ (Sup Fig 5) and went on to perform synergistic studies using IC90, IC45 and IC22.5 of Molnupiravir, Nirmatrelvir and Remdesivir in combination with IC45 and IC22.5 of Auranofin. Remdesivir alone at IC90 achieved approximately a 73% reduction in infection, but when combined with Auranofin at IC45, the reduction increased to 86%, which is a 13% increase in antiviral efficacy (Fig 5A). A similar 13% increase in effectiveness was observed when cells were treated with both Remdesivir and Auranofin at IC45 each. Individually, these doses reduced infection by about 64%, whereas their combination led to a 77% reduction (Fig 5 A,B). Although similar drug combinations with Molnupiravir and Auranofin also resulted in more inhibition compared to individual drugs, the overall synergy score was less (synergy score 1.96) (Fig 5 C, D). Nirmatrelvir and Auranofin combination showed the lowest synergy score of -5.89 (Fig 4 E, F). Overall, the Auranofin and Remdesivir combination showed the best synergistic antiviral effects with the highest synergy score of 10.19 (Fig 5 B). To better recapitulate the synergistic antiviral effects with Remdesivir in animal models, we employed GS-621763, an orally bioavailable prodrug of the remdesivir parent nucleoside GS-441524.^35^ We first evaluated the combinatorial activity in vitro using HEK293T ACE2 cells infected with SARS-CoV-2 Hong Kong strain. Co-treatment with sub-IC_50_ concentrations of Auranofin and GS-621763 caused a 3-log10 reduction of vRNA load compared to virus control (Sup Fig 6 A). This was 2-log magnitude lower than the condition with Auranofin alone, which was also evident by >1 log10 reduction in infectious virus load (Sup Fig 6 A,B). A similar trend was seen when the drugs were tested against Delta variant, albeit at a comparatively reduced magnitude, compared to SARS-CoV-2 Hong Kong strain (Sup Fig 6 C). Notably, cotreatment led to the complete loss of detectable infectious viruses in cell culture supernatants (Sup Fig 6 D). The residual RNA signal detected by qRT-PCR likely reflects the presence of non-infectious viral genomes or genome fragments that might persist despite the absence of replication-competent viruses. Upon testing the effect of the drugs against the Omicron variant, Auranofin exhibited only modest antiviral activity on its own, while GS-621763 alone and combination treatment completely abrogated vRNA and infectious viral loads which indicates a greater susceptibility of Omicron to the combinatorial treatment (Sup Fig. 6E,F).

**Fig 5.**
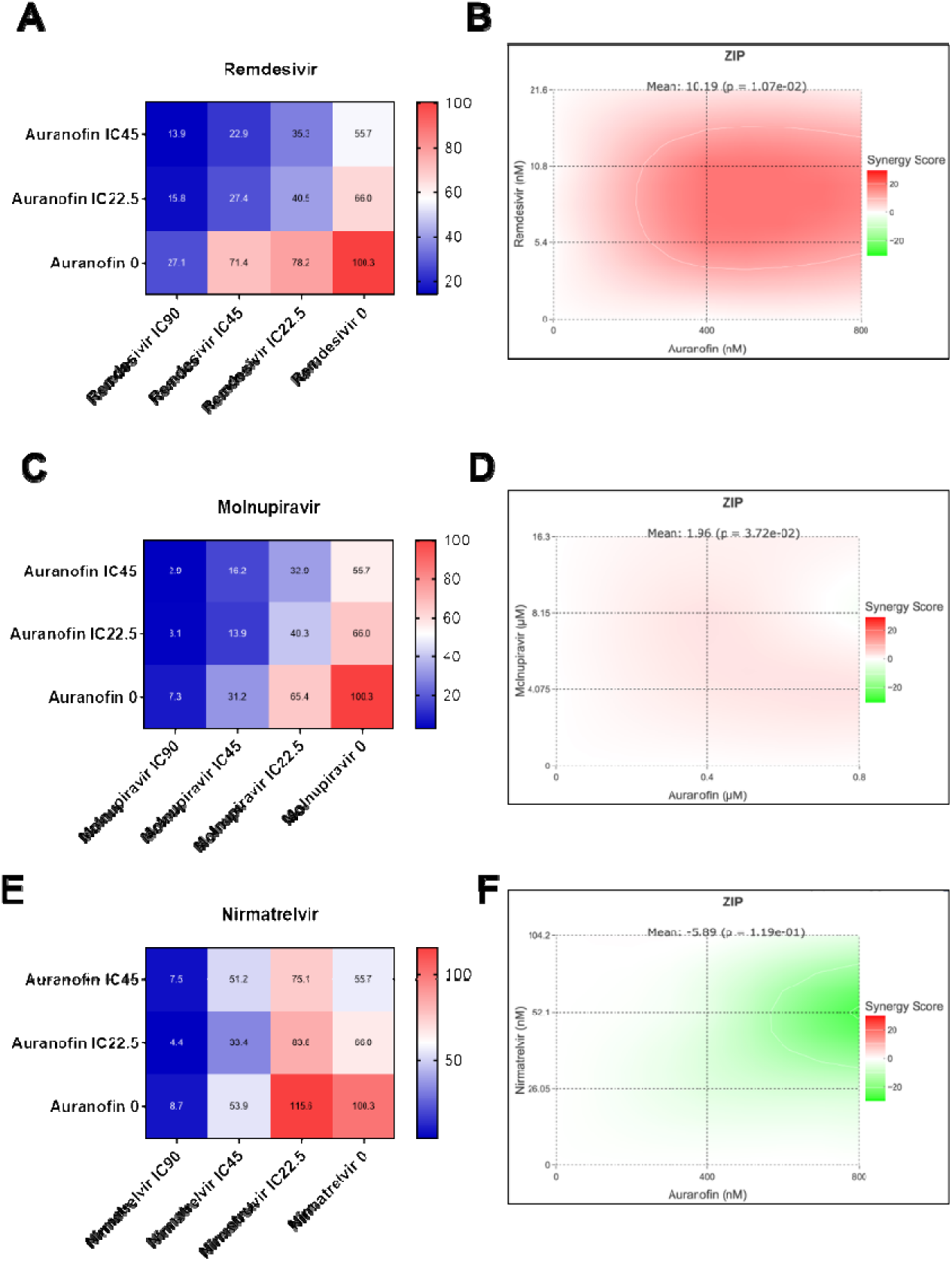
SARS-CoV-2 Antiviral efficacy of Auranofin in combination with DAAs in cell culture model. HEK293T ACE2 cells were pre-treated for 3 h with indicated concentrations of Auranofin in combination with Molnupiravir, Nirmatrelvir, or Remdesivir. Cells were then infected with 0.1 MOI SARS-CoV-2 Luc in the presence of the above drug combinations and 48 hpi, cell lysates were harvested for detection of luciferase activity. Heat maps represent the percentage of virus infection corresponding to drug combinations of Auranofin with (A) Molnupiravir, (C) Nirmatrelvir, and (E) Remdesivir. (B, D, and F) Show synergy score calculated using the SynergyFinder web tool using the Zero Interaction Potency (ZIP) model based on the percentage inhibition values calculated in A, C, and E with Graph pad PRISM19. Data represents results from 3 independent replicates.

**Figure 6:**
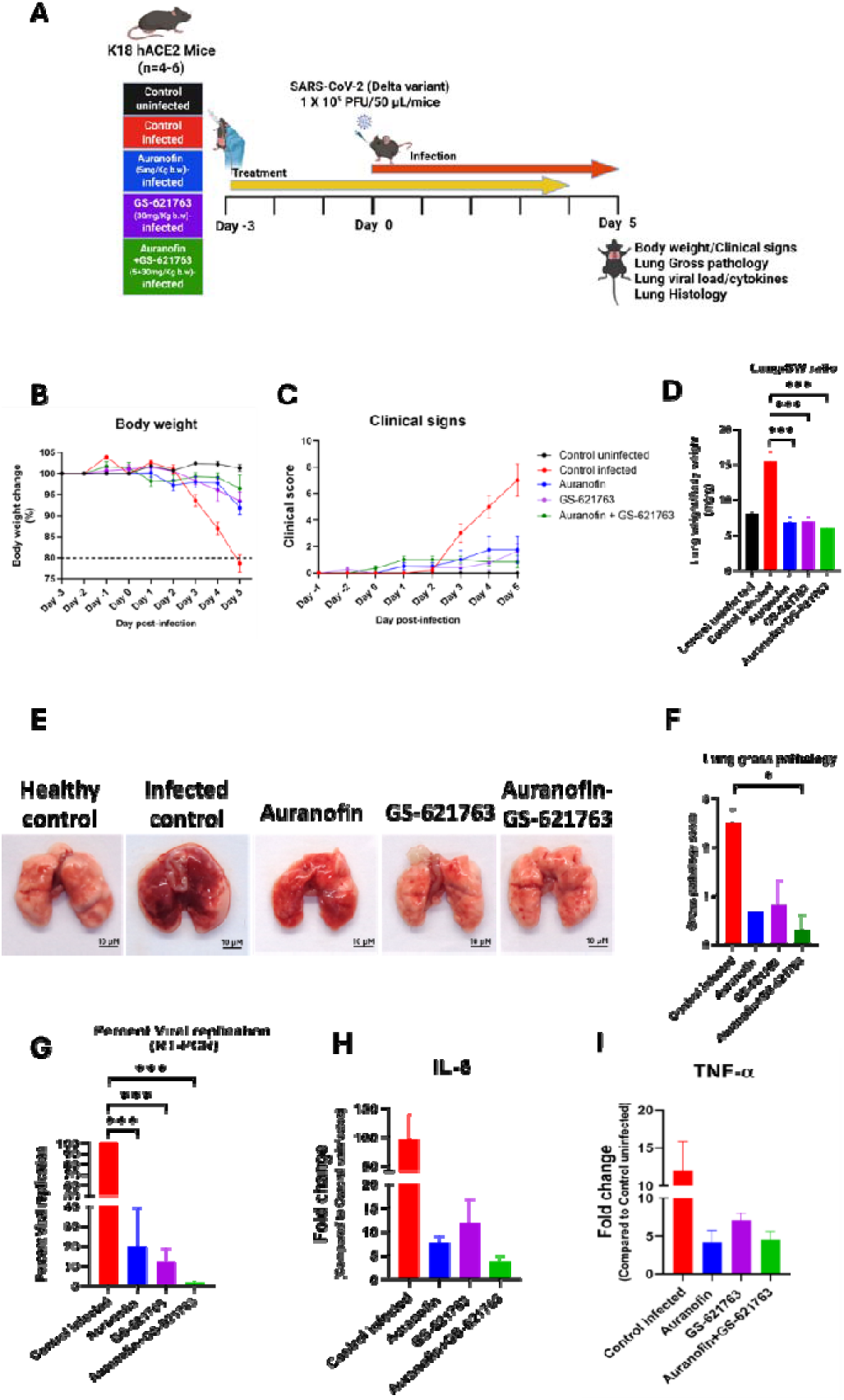
Auranofin and Remdesivir GS-621763 combination therapy shows synergistic antiviral activity in mice model. (A) Schematic showing drug dosage and infection schedule followed as part of the study. Treatment groups included (1) 5 mg/Kg b.w Auranofin, (2) 30 mg/Kg b.w GS-627163, or (3) a combination of both drugs at the same concentration. The animals were orally administered with drugs starting from D3 prior to infection, infected at D0 with 10^5^ PFU SARS-CoV-2 Delta variant, and treatment was continued until 4 dpi. Animals were sacrificed the next day, and lung tissues were harvested for further processing. BW loss and clinical signs are plotted in (B) and (C) respectively. (D-F) The lung to BW ratio comparing each group against infected control, is shown in (D), representative images of gross lung pathology from each group and corresponding pathology scoring are shown in (E) and (F) respectively. SARS-CoV-2 vRNA load quantified by qRT-PCR is shown in (G). (H,I) show IL-6 and TNF-α mRNA levels in lung homogenates quantified by qRT-PCR and represented as fold change relative to un-infected control. Data are from 1 experiment. Body weight and Clinical signs data were analysed using two-way ANOVA, Lung gross pathology, Lung-BW ratio and viral copy number data were analysed using One-way ANOVA. Statistical analyses were performed using GraphPad Prism version 8.4.3. p-values <0.05 were considered statistically significant.

### Auranofin potentiates the antiviral efficacy of GS-621763 in SARS-CoV-2 infected K18-hACE2 mice

To evaluate whether Auranofin enhances the antiviral efficacy of the orally bioavailable remdesivir prodrug GS-621763, K18-hACE2 mice were treated prophylactically beginning 3 days prior to infection with SARS-CoV-2 Delta variant and continued until 4 dpi (Fig. 6A). Vehicle-treated infected mice exhibited progressive body-weight loss following infection (Fig. 6B), accompanied by worsening clinical scores (Fig. 6C), consistent with severe disease in this model. Auranofin and GS-621763 treated mice partially mitigated body-weight loss and clinical severity (Fig. 6B,C). In contrast, combination therapy with Auranofin and GS-621763 resulted in relatively iMproved body-weight maintenance and reduced clinical scores, indicating better protection compared to either drug individually. At 5 dpi, vehicle-treated infected mice displayed significantly increased lung-to-body-weight ratios, reflecting severe pulmonary involvement such as edema (Fig. 6D), along with pronounced gross lung pathology (Fig. 6E, F). The drug treated mice showed significant (p<0.001) recovery from infection induced lung-to-body-weight ratio which was comparable to uninfected mice (Fig. 6D). Notably, treatment with Auranofin and GS-621763 alone showed a visible reduction in gross pathology, whereas combined treatment of Auranofin and GS-621763 significantly reduced gross pathology, demonstrating enhanced protection against SARS-CoV-2–induced lung injury which was aligning with both lung-to-body-weight ratios (Fig. 6D-F). Quantification of viral RNA in lung homogenates revealed high levels of SARS-CoV-2 replication in vehicle-treated infected mice (Fig. 6G). Both Auranofin (80.14 %, p<0.001) and GS-621763 (87.82 %, p<0.001) monotherapy significantly reduced lung viral RNA levels (p<0.001). Importantly, the combination of Auranofin and GS-621763 resulted in the greatest reduction in lung viral RNA (98.48 %, P<0.001), indicating enhanced antiviral efficacy relative to either agent alone (Fig. 6G).

### Combination therapy suppresses SARS-CoV-2 induced inflammatory cytokine responses

To determine whether reduced viral burden and pathology was associated with modulation of inflammatory responses, mRNA levels of IL-6 and TNF-α were assessed in lung tissues. Vehicle-treated infected mice exhibited robust upregulation of both cytokines (Fig. 6 H,I). Both Auranofin and GS-621763 treatment modestly reduced cytokine expression, while combined Auranofin and GS-621763 treatment had higher suppressed IL-6 and TNF-α expression, correlating with reduced viral replication and Improved lung pathology (Fig. 6 H,I).

## Discussion

The COVID-19 pandemic, rapid emergence of SARS-CoV-2 variants and the potential risk of antiviral drug resistance all underscore the need for therapeutic strategies that combine DAAs with HDAs. Here, we provide evidence to demonstrate that the FDA-approved anti-inflammatory drug Auranofin exerts antiviral activity against SARS-CoV-2 through a multifaceted mechanism targeting both the early and late stages of infection and exhibits robust synergy with Remdesivir and its orally bioavailable prodrug GS-621763. We followed up on our earlier report on Auranofin as a potential antiviral against SARS-CoV-2. Here, we first demonstrated that Auranofin abrogates SARS-CoV-2 entry into host cells, as shown by the SARS-CoV-2 spike pseudotype assay and ToA. These findings are consistent with a previous study suggesting that Auranofin interferes with viral entry.^36^ In addition to its effects on virus entry, we have shown that Auranofin inhibits SARS-CoV-2 Mpro activity, as demonstrated by the SARS-CoV-2 FlipGFP^Mpro^ assay. Computational docking simulations showed Auranofin binds to the SARS-CoV-2 Mpro catalytic site and forms hydrogen bonding interaction with the catalytic dyad residues. Auranofin is known to bind within the SARS-CoV-2 Mpro active-site pocket and directly interacts with key catalytic residues including Cys145 and His41.^37^ Competitive binding experiments with Auranofin and Nirmatrelvir indicates overlapping binding sites within Mpro. Hence, the relatively weak binding affinity of Auranofin suggests it may not function as a standalone Mpro inhibitor, but its dual targeting of host and virus factors likely contributes to its antiviral efficacy. Our evidence for the host-targeting effects of auranofin comes from mechanistic studies which establish that Auranofin inhibits SARS-CoV-2 entry by impairing early events during SARS-CoV-2 lifecycle, specifically post-attachment trafficking of incoming virions. Although viral binding onto the cell surface remained unaffected, Auranofin significantly reduced colocalization of virus spike with the early endosomal marker EEA1, suggesting disruption of endosomal entry or trafficking. In line with a previous report showing that Auranofin inhibits SARS-CoV-2 virus entry by perturbation of cellular redox homeostasis and lipid raft organization ^36^, our findings further demonstrate that auranofin does not induce detectable alterations in Spike stability or oligomeric state. Furthermore, Auranofin treatment did not impact SARS-CoV-2 polymerase activity, indicating no effects on viral genome transcription, translation, or replication. However, virus-induced syncytia was effectively inhibited by the drug which could be attributed to Auranofin’s inhibitory effect on virus entry either by direct effects on virus particles or host endocytic processes as reported earlier.^36^ By limiting SARS-CoV-2 spike-mediated cell fusion, Auranofin could further restrict viral dissemination, in addition to its effects on viral replication. Auranofin’s known anti-inflammatory effects ^36,38^ are supported by our findings showing inhibition of the NFκB pathway both in response to TNF-α stimulation as well as SARS-CoV-2 infection. These results underscore Auranofin’s dual action both as an antiviral agent and as an anti-inflammatory drug.

Among the DAAs tested, the Auranofin-remdesivir combination yielded the highest synergy scores in vitro, outperforming combinations with Molnupiravir or Nirmatrelvir. Importantly, this synergy was preserved with GS-621763 as well, across multiple SARS-CoV-2 variants. The use of GS-621763 enabled evaluation of this combination in vivo, where the combined treatment with Auranofin yielded superior protection in K18-hACE2 mice compared to monotherapy. Importantly, treatment of SARS-CoV-2 infected mice with this drug combination resulted in a near-complete suppression of viral RNA levels in lung tissues, along with improved body-weight maintenance, reduced clinical severity and diminished lung pathology. The enhanced reduction in inflammatory cytokines, especially IL-6, further supports an immunomodulatory benefit.

Overall, our study identifies Auranofin as a multifunctional antiviral agent that targets multiple phases of the SARS-CoV-2 lifecycle including entry, Mpro activity, syncytia formation, and inflammatory signalling. The dual use of auranofin and GS-621763 has very important translational value. Both drugs are orally bioavailable and auranofin has a well-established clinical safety profile, raising the possibility of using this oral combination regimen for early intervention or outpatient treatment of COVID-19, with minimal likelihood of resistance emergence.

## Materials and Methods

### Ethics Statement

All the animal procedures related to SARS-CoV-2 work were approved by the Institutional Animal Ethics Committee and were performed in a Viral Biosafety Level-3 facility, Indian Institute of Science, Bangalore.

### Drugs

Auranofin (Cat No. 15316, Cayman Chemicals) was solubilized in 2.5% Dimethyl sulfoxide (DMSO) and 97.5% water, whereas GS-627163 was solubilized in a solvent mixture containing 2.5% DMSO (D2650, Sigma-Aldrich), 10% Kolliphor (07076, Sigma-Aldrich), 10% Labrasol® (Gattefosse), 2.5% Kollisolv® Polypropylene Glycol 400 (06855, Sigma-Aldrich), 75% water.

### Cells and Virus

The following cell lines were used in this study: HEK293T (NCCS, Pune, India), HEK293T cells expressing human ACE2 (HEK293T-ACE2) (NR-52511, BEI Resources), Vero E6 (CRL-1586, ATCC). All cell lines were cultured in complete Dulbecco’s modified Eagle Medium (12100-038, Gibco) with 10% HI-FBS (16140-071, Gibco), 100 IU/mL Penicillin and 100 mg/mL Streptomycin (15140122, Gibco) supplemented with GlutaMAX (35050-061, Gibco).

The following SARS-CoV-2 isolates were procured from BEI Resources, NIAID, NIH: Isolate Hong Kong/VM20001061/2020, NR52282; NR-55282, Isolate hCoV-19/USA/PHC658/2021 (Lineage B.1.617.2; Delta Variant); Isolate hCoV-19/USA/MD-HP20874/2021 (Lineage B.1.1.529; Omicron Variant), NR-56461. The SARS-CoV-2 Delta variant used for animal experiments was isolated from a human clinical sample obtained at the COVID-19 diagnostic facility in Centre for Infectious Disease Research, Indian Institute of Science, in accordance with institutional biosafety guidelines. SARS-Related Coronavirus 2, Isolate USAWA1/2020, Recombinant Infectious Clone with Nanoluciferase Gene (icSARS-CoV-2-nLuc), Catalog No. NR-54003. Both viruses were propagated and titrated by plaque assay in Vero E6 cells as previously described ^39^.

### Spike Pseudotyped virus entry assay

Pseudotyped virus particles expressing SARS-CoV-2 spike protein were prepared as described previously ^21^. HEK293T ACE2 cells were seeded in poly-L-lysine (Sigma-Aldrich, P9155-5MG) pre-coated 96-well plates and used for the experiments upon reaching 70-80% confluency. Cells were pre-treated with 1 μM Auranofin (Sigma Aldrich, A6733) for 3 h, followed by infection with pseudotyped particles in the presence of 10 μg/ml polybrene (Merck, TR-1003-G). Auranofin was present in the media throughout the experiment. After 24 h, luciferase activity was quantified with a Firefly luciferase assay kit (Promega, E4550) using a TECAN multi-mode plate reader.

### Time of addition assay

HEK-ACE2 cells were seeded in poly-L-lysine pre-coated 24-well plates to reach ∼90% confluency after 24 h. To test the effect on virus entry, cells were pre-treated with 1 μM Auranofin for 3 h and then infected with SARS-CoV-2 virus at 10 MOI. After 3 h incubation, cells were lysed, and the expression of SARS-CoV-2 spike was analyzed by western blotting. Auranofin was present in the media throughout the experiment. Quantification of band intensity was done using imagej/Fiji.

### Virus binding assay

For the SARS-CoV-2 binding assay, HEK293T ACE2 cells were trypsinised to make a cell suspension and resuspended in complete media. Cells were then infected with 50 MOI SARS-CoV-2 for 45 min on ice in the presence of 20 µM FNDR 11124. Unbound virus particles from cell surface were removed by washing twice with ice cold PBS and cells were fixed using 4% PFA for 10 min. Cells were then washed with PBS and incubated in blocking buffer (PBS containing 2% BSA) for 30 min. Detection of cell surface-bound SARS-CoV-2 spike was done by incubating the cell suspension in mouse anti-Spike primary antibody (Genetex, GTX632604) diluted in blocking buffer for 30 min. Cells were then washed with PBS and resuspended in blocking buffer containing anti-mouse Alexa 488 secondary antibody (Invitrogen, A-11001) for 30 min. After further washing, the cell population positive for spike surface staining was analysed using a Beckman Coulter CytoFLEX flow cytometer. Data was analysed using CytExpert 2.5.0.77 software with unstained and single-stained controls.

### Replicon assay

The protocol for SARS-CoV-2 replicon assay was as published before ^26^, with some modifications. Briefly, HEK293T ACE2 cells in a 24-well dish were co-transfected with 250 ng each of SARS-CoV-2 F-luc and RdRp plasmids (kind gift from Subhash C Verma)^26^ and after 6h, media was replaced with that containing 20µM FNDR 11124. 10 µM Remdesivir was used as a positive control. In both cases, cell lysates were collected after 24 h using passive lysis buffer and firefly/renilla luciferase expression were quantified using Dual-Luciferase Reporter Assay System (Promega, E1980) as per manufacturer’s instructions. Luminescence readings were taken using a Thermo Fisher Varioskan LUX multimode plate reader.

### SARS-CoV-2 FlipGFP^Mpro^ reporter assay

HEK293T cells were seeded in poly-L-lysine pre-coated 24-well plates and 24 h later used for the experiment. For optimizing the reporter system, cells were cotransfected with 125 ng of FlipGFP^Mpro^ plasmid (Addgene, 163078) and 375 ng of NSP5 WT or NSP5 C145A plasmids (kind gift from Prof. Nevan Krogan - University of California San Francisco) ^40^ using Lipofectamine 2000 transfection reagent (Invitrogen, 11668019). After 24 h, cells were fixed and GFP and mCherry expressions were analyzed under an EVOS M-5000 fluorescence microscope. For testing the effect of Auranofin, HEK293T cells were transfected with 125 ng of Mpro reporter and 375 ng of NSP5 WT plasmids, and 3 h later, media was replaced with fresh DMEM containing 1 μM Auranofin or 7.8 μM Nirmatrelvir. 36 h post-transfection, cells were fixed using 4% paraformaldehyde (PFA), and GFP fluorescence signals were analysed using ImageJ/Fiji.

### Polymerase assay

To study the effect of Auranofin on virus polymerase activity, cells were infected with 10 MOI SARS-CoV-2, and 6 h later, 1 μM Auranofin was added to the medium. Cells were further incubated for 3 h, and cell lysates were collected for western blot analysis, and total RNA was isolated using TRIzol for RT PCR.

### Syncytia assay

HEK293T cells were co-transfected with 1 μg of spike and 1 μg of GFP1-10 plasmid, and HEK293T ACE2 cells were transfected with GFP11. After 24 h, HEK293T cells were pre-treated with 1 μM Auranofin for 3 h. Transfected cells were trypsinized, and 0.1 million HEK293T cells were co-cultured with 0.1 million HEK293T ACE2 cells in the presence of Auranofin. 24 h later, cells were fixed in 4% formaldehyde, and the nucleus was stained with DAPI. Representative images were acquired in an Evos M-5000 fluorescence microscope. Total GFP intensity was measured using a CX4 High Content Screener.

### NFκB assay

HEK293T cells were seeded into poly-L-lysine coated 24-well plates and allowed to settle for 24 h before transfection. Cells were co-transfected with 50 ng of IgK-IFN-luc plasmid expressing a firefly luciferase gene driven by the NFκB promoter (Addgene, 14886), 20 ng of pRL-TK plasmid expressing constitutively active renilla luciferase (provided by Prof. Adolfo García-Sastre (Icahn School of Medicine at Mount Sinai, New York) and has been described before ^41^, and 500 ng of empty vector. After 24 h, cells were pre-treated with 1 μM Auranofin for 3 h and stimulated with 50 ng/well of TNF alpha (PeproTech, 300-01A-100 UG) for 12 h. Luciferase activity was quantified with a dual-luciferase kit (Promega Cat# E1980) as per the manufacturer’s instructions, using a TECAN Infinite 200-PRO multiplex reader. To evaluate the effect of Auranofin on the NFκB pathway during viral infection, cells were transfected with reporter plasmids as above, but stimulated by infecting with 0.1 MOI SARS-CoV-2 for 48 h. Dual luciferase activity was measured as above. Human TNF alpha expression levels in infected cells were quantified by qRT PCR using the following primers: Forward primer: 5’ AGTGAAGTGCTGGCAACCAC 3’ and Reverse primer: 5’ GAGGAAGGCCTAAGGTCCAC 3’.

### In Silico Drug Docking Analysis and Structure visualisation

The 3D structure of Auranofin was built and optimized using Avogadro. The structure of Nirmatrelvir was obtained from DrugBank, and the SARS-CoV-2 Mpro structure (PDB: 8DZ2) was retrieved from the Protein Data Bank. The Mpro structure was processed using Autodock 4.2.6, which included repairing missing atoms, removing water molecules, adding hydrogen atoms, and assigning Kollman charges. Similarly, the structure of Auranofin and Nirmatrelvir compounds was processed by assigning Gasteiger partial charges. The grid box was centered on the Mpro active site with a spacing of 0.297 Å to ensure complete coverage of the binding region. Docking simulations were performed using 200 genetic algorithm runs, with a population size of 500, 27 million energy evaluations, and 100,000 generations. Individual docking poses were clustered using a root-mean-square deviation (RMSD) cutoff of ≤2.0 Å. The cluster with the most members and the lowest binding free energy was selected as the preferred docking pose. The best-docked poses were visualized and analysed using ChimeraX, where hydrogen-bonding interactions between drugs and Mpro residues were identified and examined.

### Isothermal Titration Calorimetry for Auranofin and Nirmatrelvir with SARS-CoV-2 Mpro

Isothermal Titration Calorimetry (ITC) (MicroCal ITC200, Malvern, Northampton, MA) was used in this experiment. The binding isotherms obtained from ITC experiments were analyzed using the non-linear binding model provided by Origin software (MicroCal, Inc.). SARS-CoV-2 Mpro was purified via Ni-NTA affinity chromatography as previously ^42^. The purified Mpro was dialyzed at 4 °C in buffer A (20 mM HEPES, 50 mM NaCl, pH 7.5) for ITC experiments. Stock solutions of both compounds (10 mM) were prepared in DMSO and stored at -20 °C. The final DMSO concentration during experiments was maintained at 2% in both the cell and syringe samples to eliminate any heat change generated due to DMSO. Prior to the experiments, both the protein sample and the compounds were degassed to prevent bubble formation.

For the ITC experiments, Mpro (20 μM) in buffer A was loaded into the titration cell and 200 μM final concentration of the compounds prepared in buffer A were placed in the injection syringe. The final concentration of DMSO used during the experiments were 2%. The titration involved 20 injections: one initial injection of 0.4 μL followed by 19 injections of 2 μL each, with a 220-second interval between injections. The system temperature was maintained at 25 °C, and the reference power was set to 8 μcal/s. After obtaining the binding isotherms for both compounds, sequential binding experiments were performed to examine how one compound influences the binding of the other. For these experiments, Mpro (20 μM) was preincubated with 200 μM Auranofin at 4 °C for 2 hours and loaded into the titration cell. Nirmatrelvir (200 μM) was titrated against the preincubated Mpro-Auranofin complex under identical experimental conditions. A reverse experiment, where Auranofin was titrated against the Mpro-Nirmatrelvir complex, was also conducted. The resulting thermograms were analyzed using the Origin software package to determine binding parameters, including the association constant (Ka = 1/Kd), enthalpy (ΔH), and entropy (ΔS).

### IC90 calculation of the drugs

HEK293T ACE2 cells were seeded in poly-L-lysine pre-coated 24-well plates and 24 h later cells were pre-treated for 3 h with increasing concentrations of Molnupiravir (BLD pharmatech, BD01383150) (20, 10, 5, 2.5 and 1.25 μM), Nirmatrelvir (Medchem express, HY-138687) (100, 80, 75, 60, 50, 25, 16, 10 nM) or Remdesivir (Cat. no. 30354 Caymen Chemicals) (1 μM, 0.5 μM, 0.25 μM, 125 nM, 100 nM, 60 nM, 20 nM, 16 nM, and 4 nM). Cells were then infected with SARS-CoV-2 Luc at 0.1 MOI. After 48 h, luciferase activity was measured using the Promega E2820 Renilla Luciferase Assay System as per the manufacturer’s instructions. Luminescence measurements were taken using a TECAN Infinite 200-PRO multiplex reader. Percentage infection was calculated and normalized to untreated control and IC90 was determined in GraphPad Prism.

### Antiviral testing of Auranofin in combination with DAAs

HEK293T ACE2 cells were seeded in poly-L-lysine pre-coated 24-well plates and 24 h later cells were pre-treated for 3 h with IC90, IC45, or IC22.5 of Molnupiravir, Nirmatrelvir, Remdesivir with IC45 or IC22.5 of Auranofin. Cells were then infected with 0.1 MOI SARS-CoV-2 Luc in the presence of the above-mentioned drug combinations and after 48 h, luciferase activity was measured using measured using Promega E2820 Renilla Luciferase Assay System as per manufacturer’s instructions. Luminescence measurements were taken using a TECAN Infinite 200-PRO multiplex reader. The synergy scores were determined using the SynergyFinder web tool ^43^ employing the Zero Interaction Potency (ZIP) model ^44^.

### Statistics & Illustrations

All statistical analyses used in this manuscript were performed using GraphPad Prism 8.4.3 (GraphPad Software, USA). Information on the statistical method used is mentioned in the legend section of the respective figures. Error bars indicate SD. In all cases, a p-value <0.05 is considered significant. Figure illustrations have been generated using Biorender.com.

### SARS-CoV-2 infection experiments in animal model Animals, treatment regime and infection

Transgenic K18-hACE2 mice (initial breeders obtained from The Jackson Laboratory (Strain: 034860, USA) was maintained and bred in the institutional central animal facility under controlled temperature, humidity and a 12h light-dark cycle with ad libitum access to standard feed and water. Transgenic K18-hACE2 Mice (n=4-6) of 8-10 weeks old were allocated randomly into five groups. For treatment groups, mice were administered orally twice a day with 200µL of Auranofin (5mg/Kg b.w.) or GS627163 (30 mg/Kg b.w.) or combination of both Auranofin (5mg/Kg b.w.) and (30 mg/Kg b.w.), beginning 3 days prior to SARS-CoV-2 infection and continued until 4 days post-infection (dpi). Control groups received the corresponding vehicle on the same schedule.

On Day 0, mice were anesthetized with Ketamine/Xyalzine mix and infected with 105 PFU of SARS-CoV-2 (Delta variant) in 50µL of PBS intranasally. Animals were monitored daily for body weight changes and clinical signs throughout the experiment. Hamsters were monitored daily for the development of clinical symptoms and scored according to predefined severity criteria. Clinical parameters evaluated included lethargy (0 = absent, 1 = mild, 2 = severe), piloerection (0 = absent, 1 = mild, 2 = moderate, 3 = severe), abdominal respiration (0 = absent, 1 = mild, 2 = severe), and hunched back(0 = absent, 1 = mild, 2 = severe). Body weight loss was also included as a clinical parameter and graded on a three-point scale based on percentage reduction relative to baseline (1-5% = 1; 5.1-10% = 2; 10.1-15% = 3). All clinical observations were recorded for each experimental group through day 4 dpi. At 5 dpi, mice were euthanized with overdose of Ketamine and lung tissues were harvested and weighed. One portion of it was used for histology and another portion was used for RNA extraction. Tissue was homogenized with a FastPrep®-24 Classic bead beating grinder (116004500, MP Biomedicals). Total RNA was extracted using RNAiso Plus (9109, TaKaRa) reagent and stored at -80°C until use.

### Quantification of Viral RNA by RT-qPCR

SARS-CoV-2 viral RNA levels in lung homogenates were measured by real-time quantitative reverse transcription PCR (RT-qPCR) targeting the Nucleocapsid-1 (N1) gene. Assay was performed using the AgPath-ID™ One-Step RT–qPCR Kit (AM1005, Applied Biosystems) in a 10 µL reaction mixture containing 100 ng of RNA per sample, run in a 384-well format plate. Comparative threshold (Ct) values were obtained and viral copy numbers were determined using a standard curve generated from SARS-CoV-2 genomic RNA. Primers and probe used are as follows: Nucleocapsid-1 forward:5′-GACCCCAAAATCAGCGAAAT-3′, reverse: 5′-TCTGGTTACTGCCAGTTGAATCTG-3′, N1 (6-FAM/BHQ-1) 5′-ACCCCGCATTACGTTTGGTGGACC-3′.

### Quantification of cytokine mRNA expression by RT-qPCR

The mRNA expression levels of TNF-α and IL-6 were quantified using extracted total RNA, with 18S rRNA serving as the internal reference gene. Complementary DNA (cDNA) was synthesized from 1 µg of RNA using the PrimeScript™ RT Reagent Kit with gDNA Eraser (RR047A, TaKaRa). RT–qPCR was performed using 2.5 µL of cDNA, 2.5 µL of gene-specific primer mix, and 5 µL of SYBR™ Green PCR Master Mix (Applied Biosystems). The primer sequences used were as follows: TNF-α for-<colcnt=4>ward: 5′-AGAAACACAAGATGCTGGGACAGT -3′, reverse:5′-CCTTTGCAGAACTCAGGAATGG-3′;IL-6 forward:5′-CTGCAAGAGACTTCCATCCAG-3′, reverse: 5′-GGTATGCTAAGGCACAGCAC AGTGGTATAGACAGGTCTGTTGG -3′. Relative gene expression was calculated using the ΔΔCt method with 18S rRNA normalization, and the mean log_2_ fold change for each gene was determined for each experimental group.

## Statistical Analysis

Body weight and Clinical signs data were analysed using Two-way ANOVA, Lung gross pathology, Lung-BW ratio and viral copy no. data were analysed using One way ANOVA. Statistical analyses were performed using GraphPad Prism (version 8.4.3). P-values <0.05 were considered statistically significant.

## Acknowledgment

This research has been supported by ICMR (IIRPIG-2023-0000978) and BIRAC grant (BT/CS0070/06/22) to ST. We acknowledge infrastructure and research support to IISc from the Crypto Relief Fund, L & T Trust, DST-FIST program, Institute of Eminence Fund, Ministry of Education, and DBT-IISc partnership program (Phase II). RN acknowledges DBT-RA fellowship, SK acknowledges PMRF fellowship and RS acknowledges FICCI-PMRF fellowship.

## List of abbreviations

SARS-CoV-2: Severe Acute Respiratory Syndrome Coronavirus-2
COVID-19: Coronavirus, COVID-19
DAA: Direct-acting antivirals, HDA, Host-directed antivirals
hpi: hours post-infection
dpi: days post-infection
HEK293T ACE2: HEK293T cells expressing human ACE2 receptor
Infection media: DMEM with 2%FBS
ToA: Time of addition
MOI: Multiplicity of Infection
ZIP: Zero Interaction Potency.

---

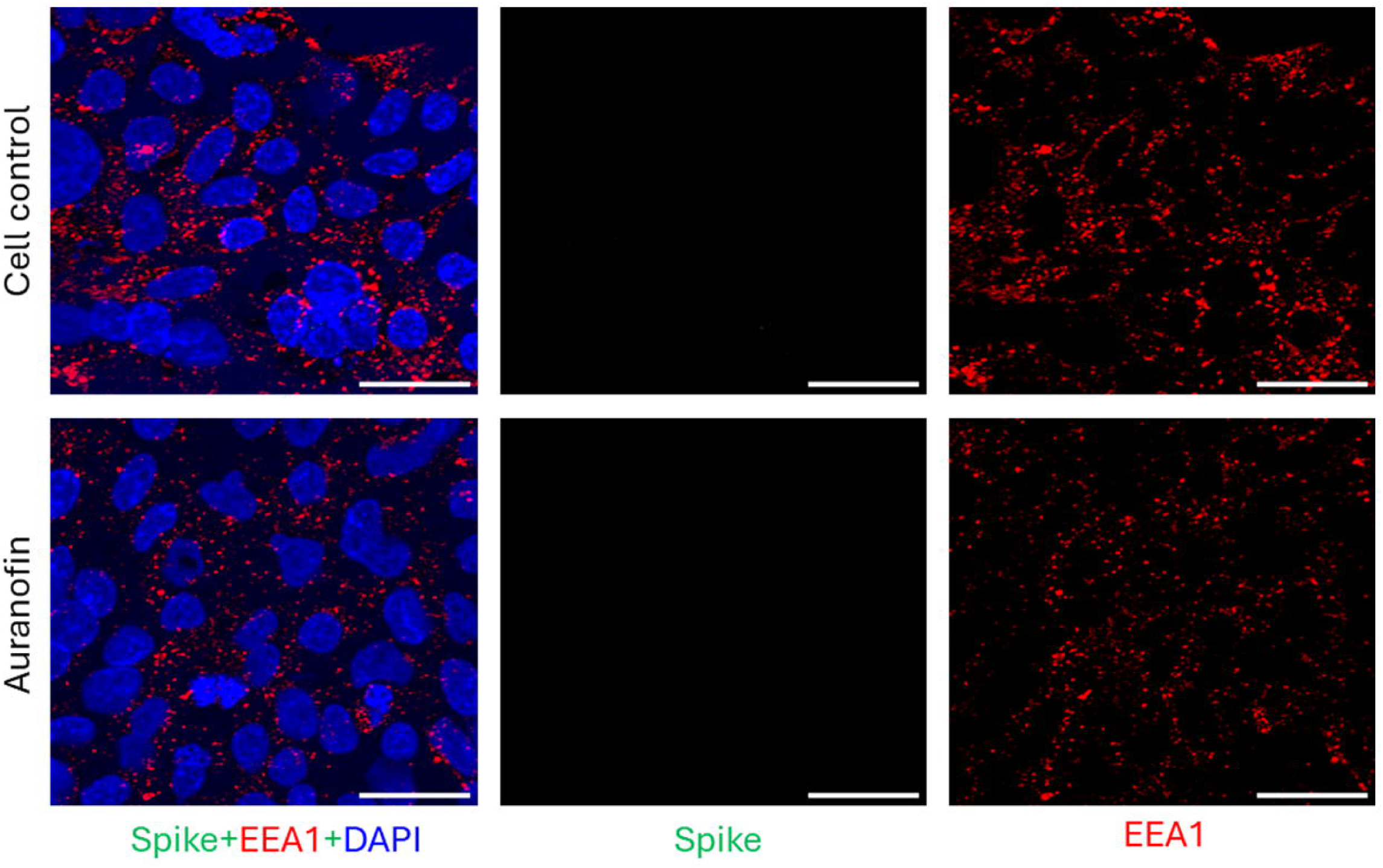

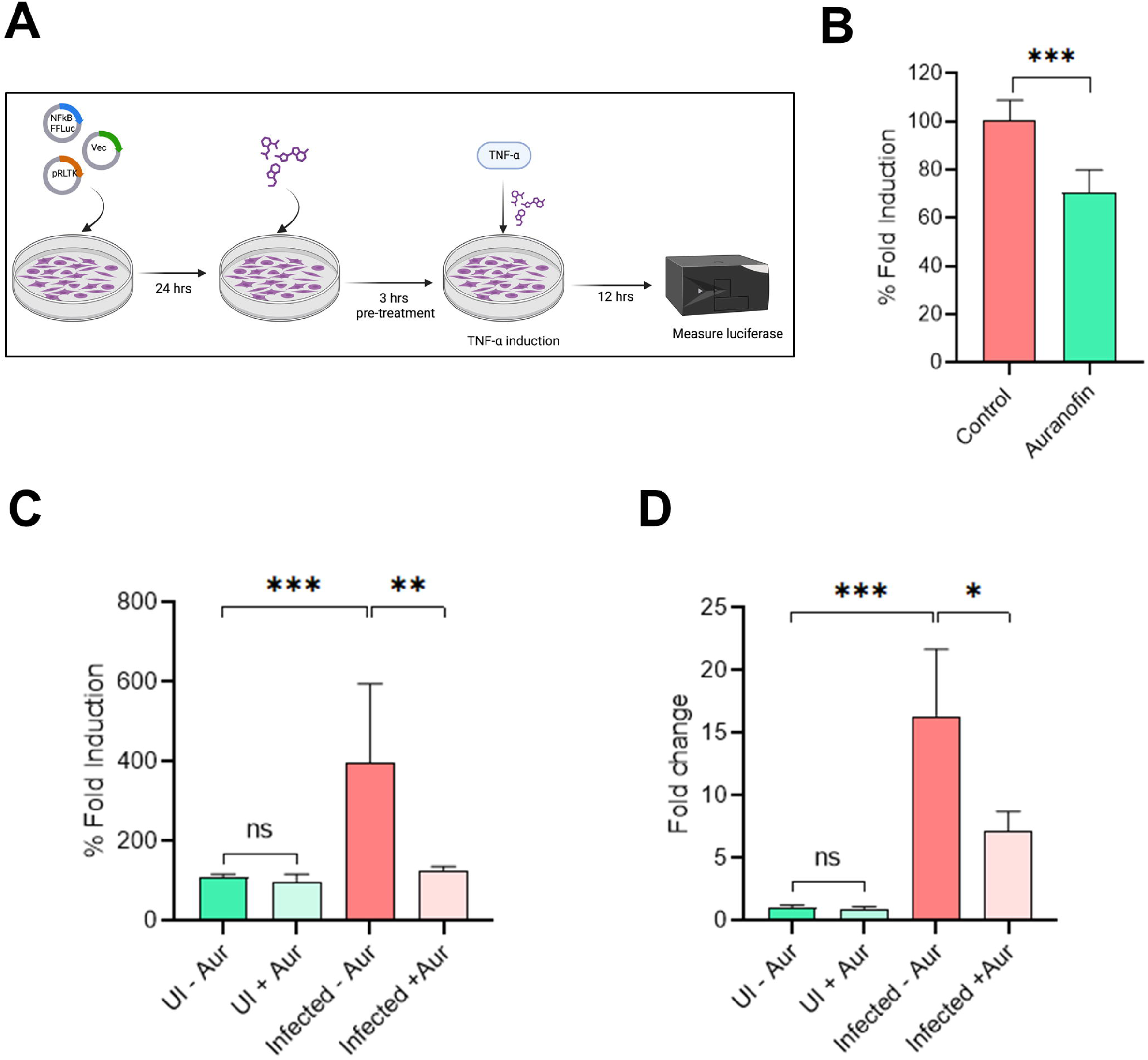

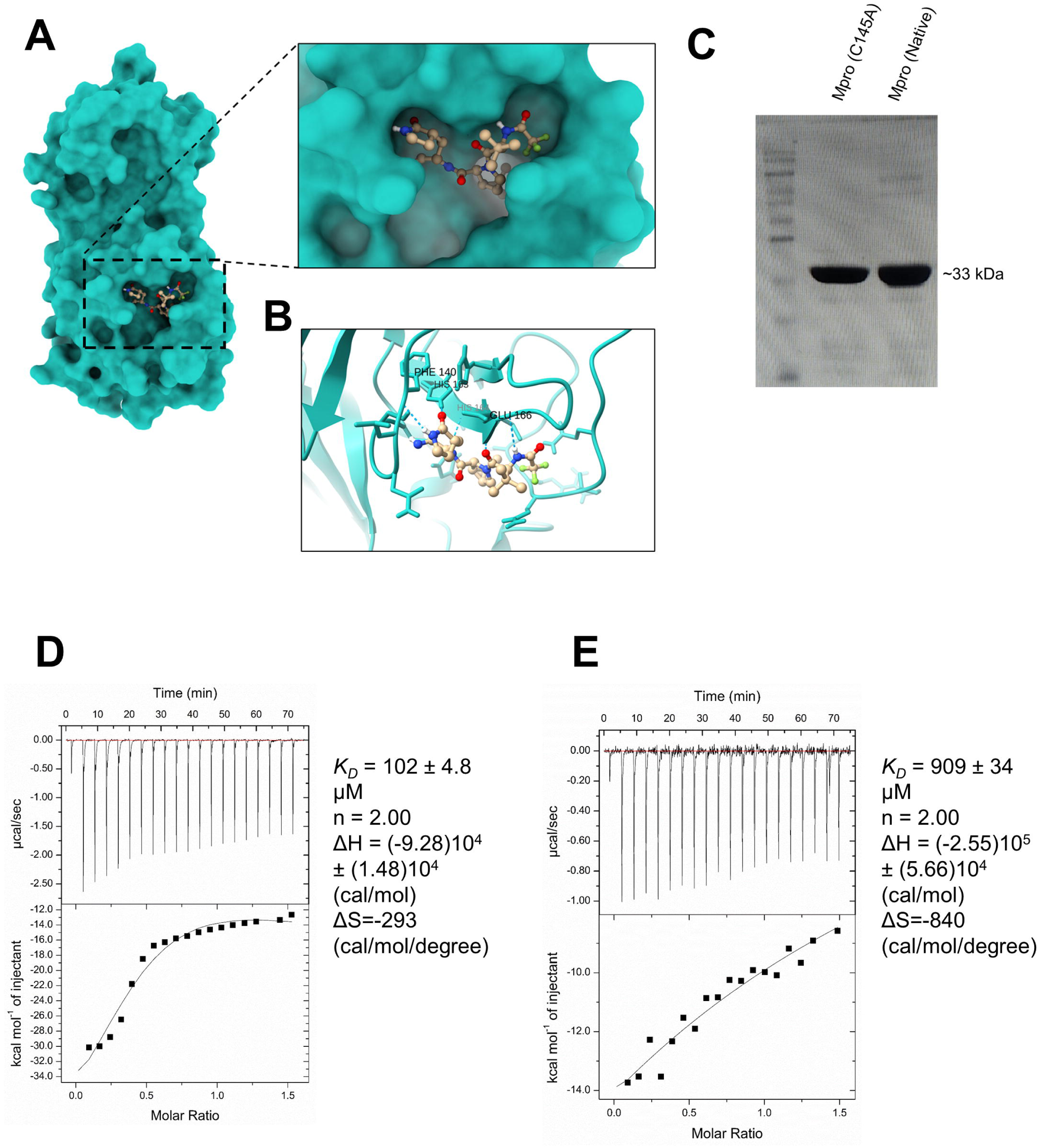

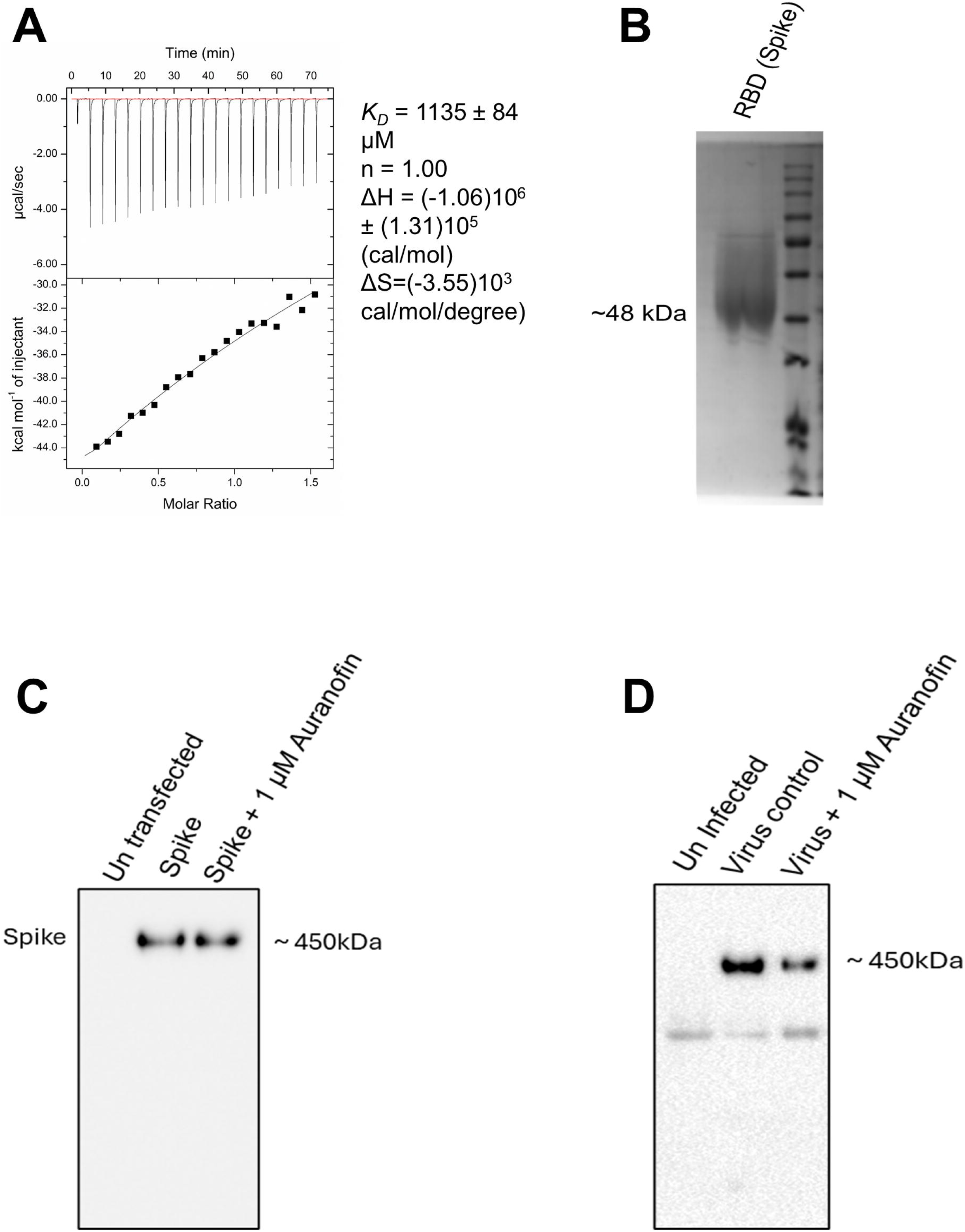

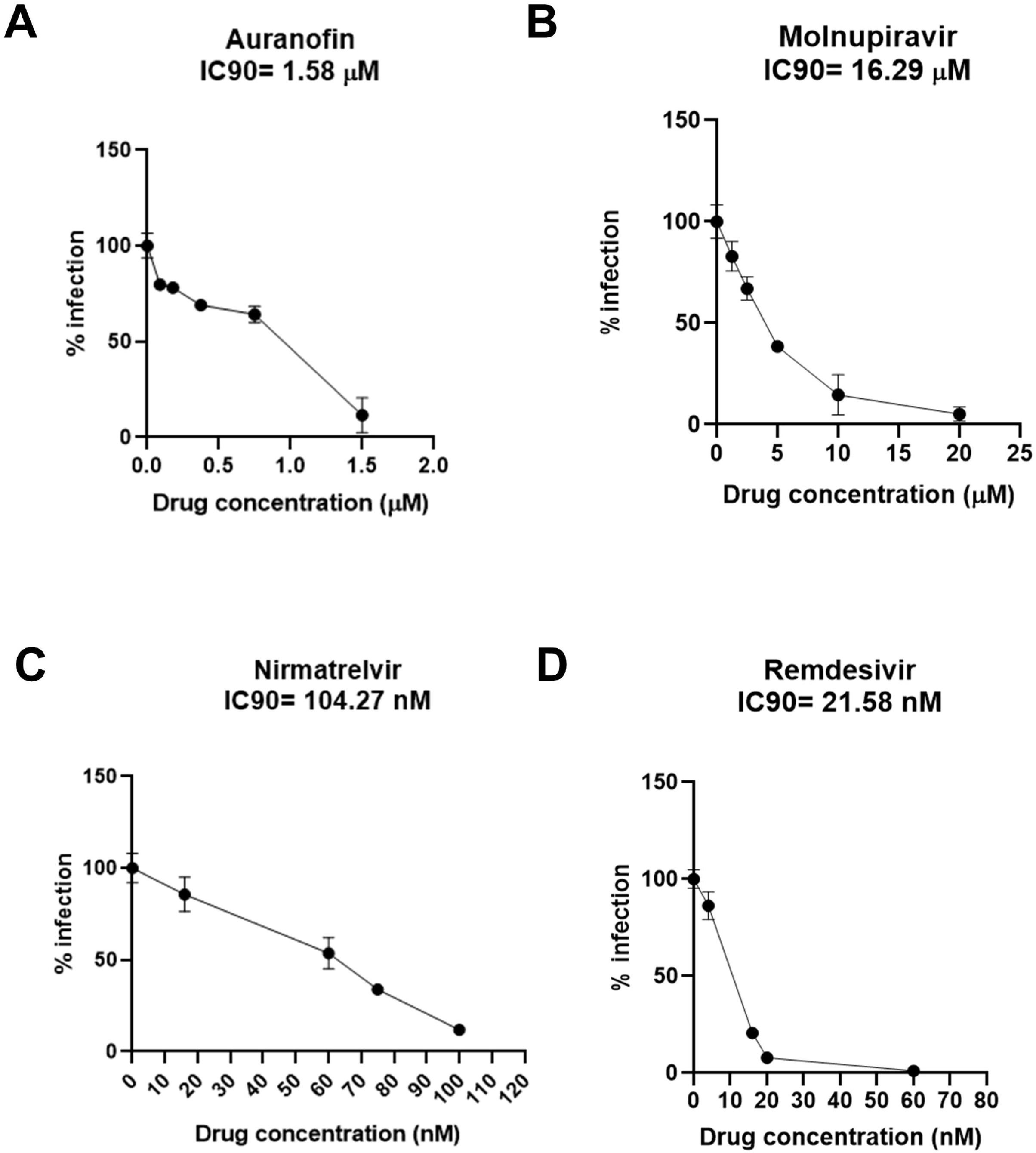

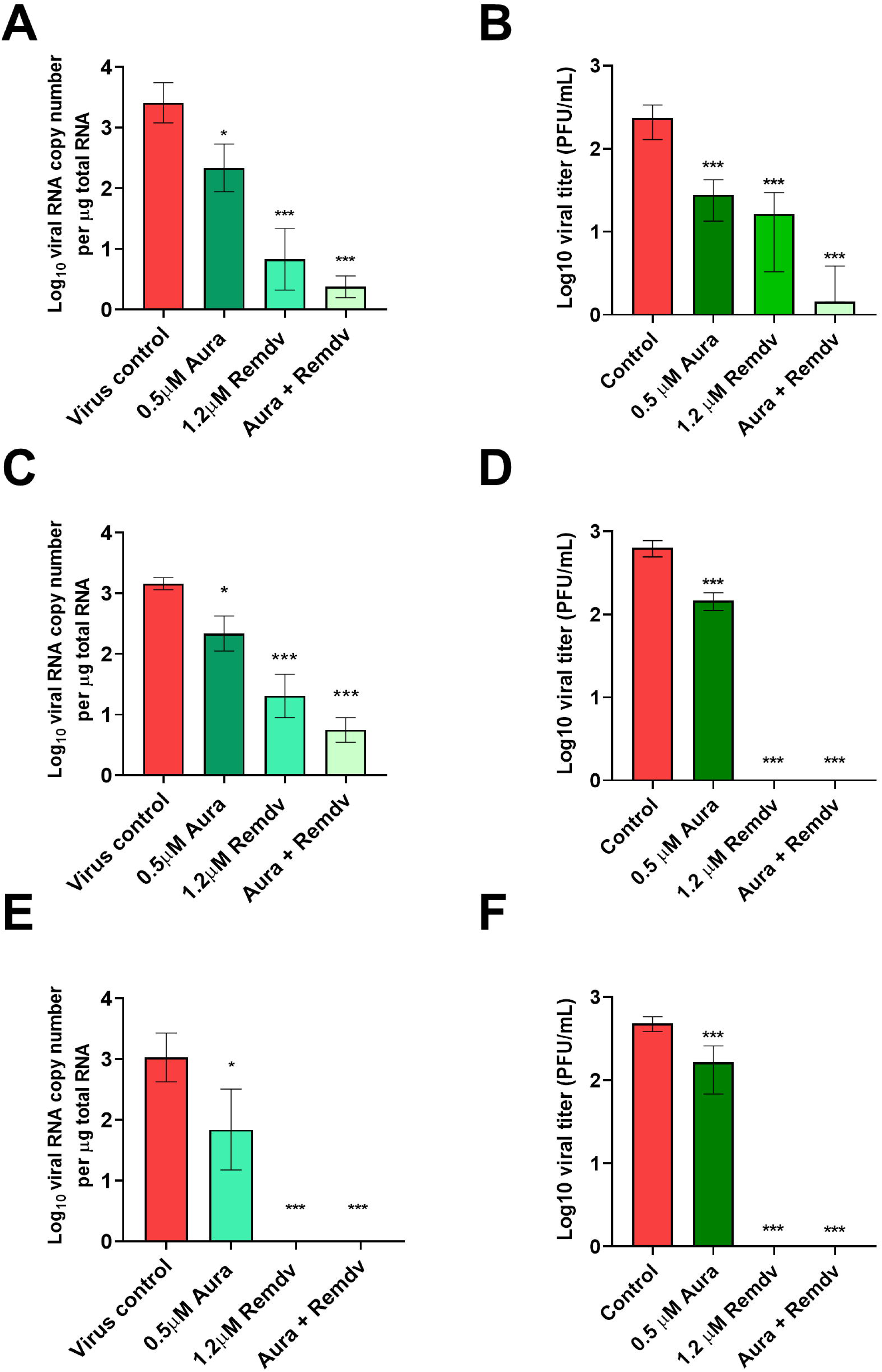

