## Supplementary data for "An Oral Combination therapy against SARS-CoV-2 based on Synergistic Action of Auranofin and Remdesivir"

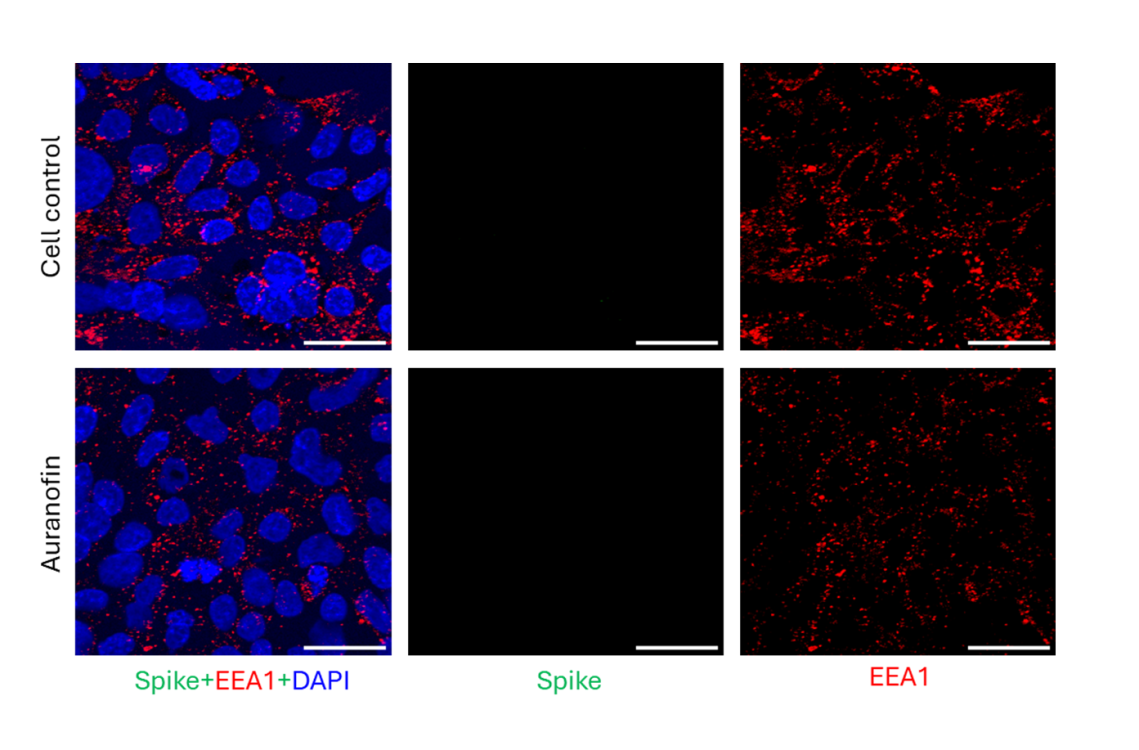

**Supplementary Fig 1**. **EEA-1 positive endosome localization in un-infected cells treated with Auranofin.** HEK293T ACE2 cells were pre-treated with 1 µM auranofin, incubated on ice for 1 h and moved to 37ºC in the presence of drug for 20 min. Cells were then fixed and labelled with antibodies against spike and EEA-1.

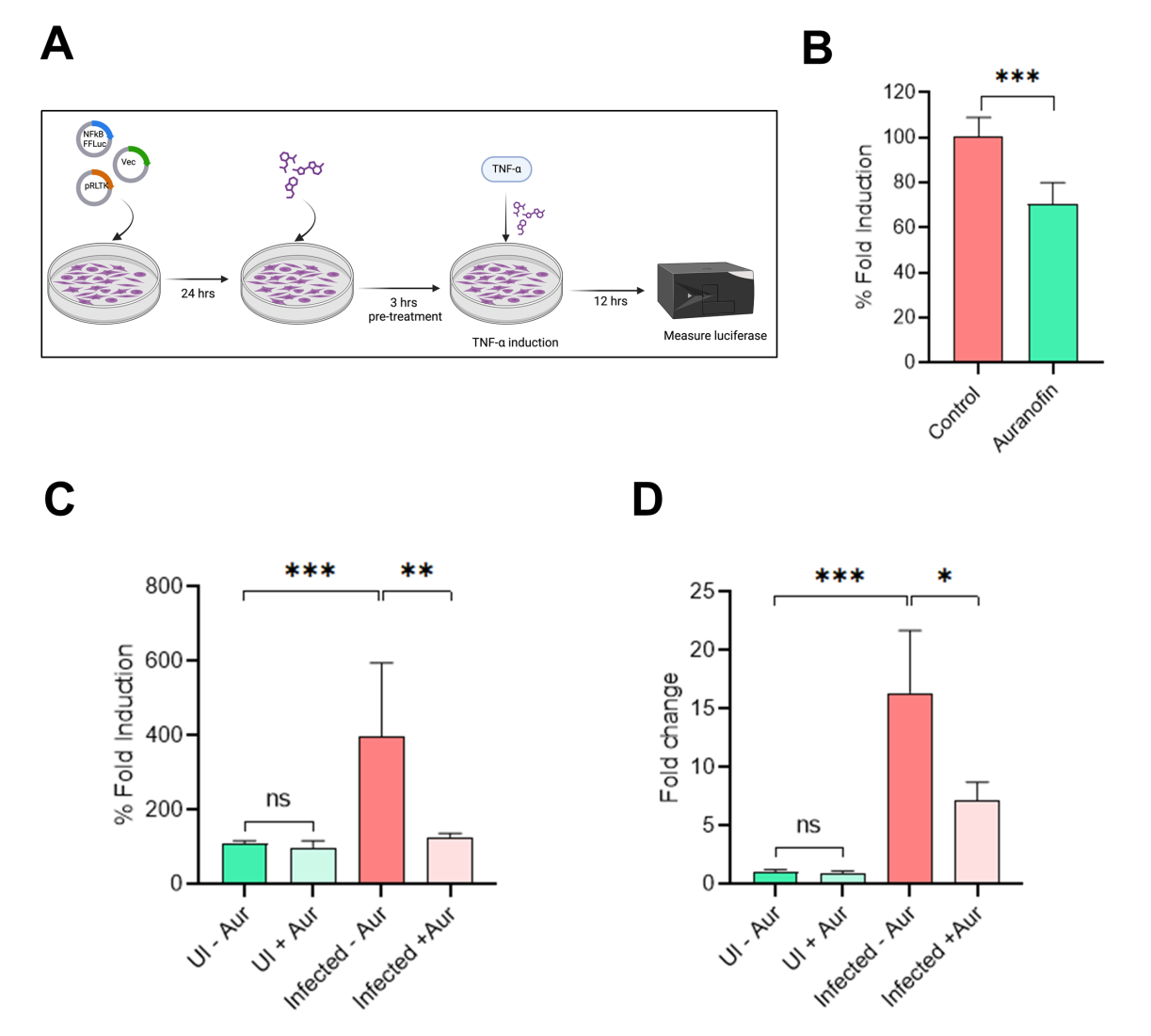

**Supplementary Fig 2. Auranofin inhibits NFκB activity and TNFa induction during SARS-CoV-2 infection**. (A) Schematic of dual luciferase assay for the analysis of NFkB activity. Briefly, HEK293T cells were co-transfected with firefly luciferase reporter plasmid driven by NFkB promoter, renilla luciferase reporter plasmid, and empty vector. After 24 h, cells were treated with 1 μM Auranofin for 3h, followed by induction of NFkB pathway using recombinant TNF alpha. Dual luciferase activity analyzed after 12 h is shown in (B). Similarly, the assay was repeated in HEK293T ACE2 cells transfected with the above plasmids but infected with 0.1 MOI SARS-CoV-2 to induce the NFkB pathway. Infected cells were collected at 48 hpi and dual luciferase assay data is shown in (C). In (B, C) luciferase activity is normalized to induced untreated control. TNF alpha expression levels in infected cells quantified by qRT PCR are shown in (D). (D, E) are from 1 experiment. (B) is from 3 independent biological replicates. ∗p < 0.05, ∗∗p < 0.01, ∗∗∗p < 0.001; using two-tailed unpaired t-test or Welch ANOVA with Dunnett’s T3 multiple comparison tests, where applicable. Error bars represent mean ± SD.

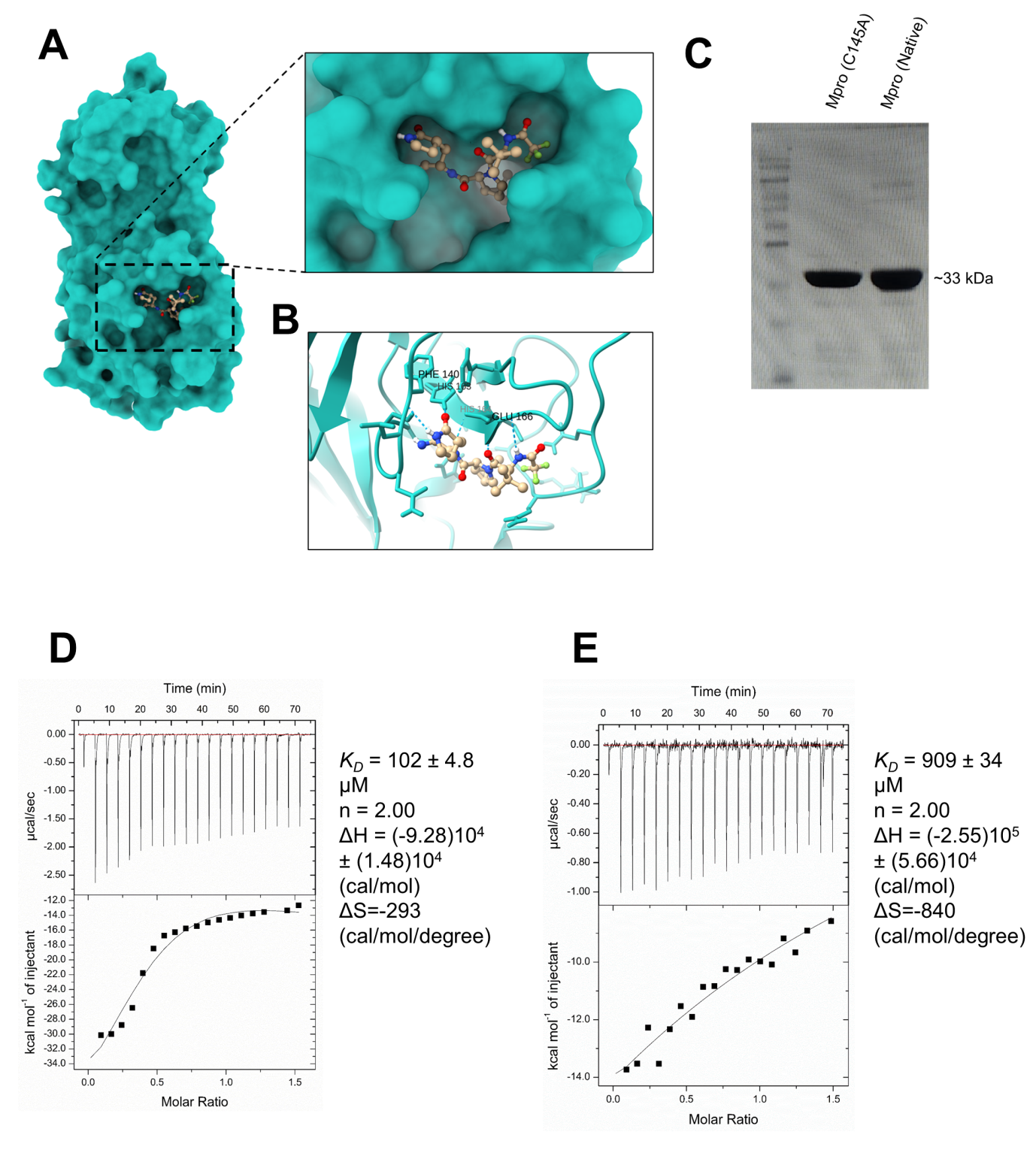

**Supplementary Fig 3. Nirmatrelvir outcompetes auranofin in binding with SARS-CoV-2 M^pro^.** Binding free energy and docking of Nirmatrelvir against SARS-CoV-2 Mpro were performed using Autodock. (A-B) Docked structures of SARS-CoV-2 Mpro to Nirmatrelvir and corresponding Hydrogen bonding interactions with key amino acids are shown in A and B respectively. (C) Western blot data for mutant main protease (Mpro, C145A), and native Mpro. (D,E) ITC Sensogram and isothermodynamics for (D) Nirmatrelvir titrated against Auranofin-M^pro^ complex and (E) auranofin titrated against nirmatrelvir-Mpro complex.

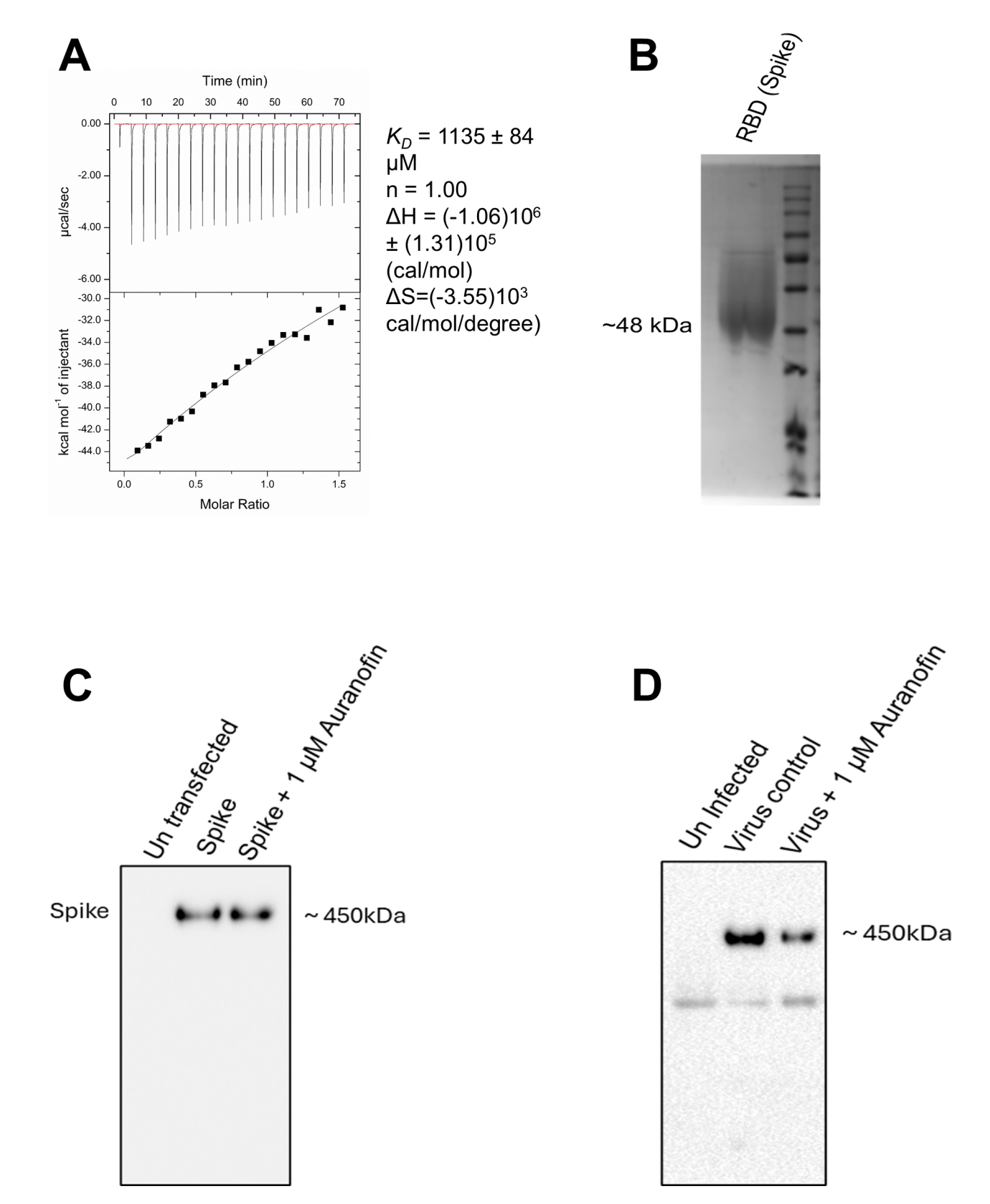

**Supplementary Fig 4. Auranofin does not induce gross structural alterations in the SARS-CoV-2 spike protein.** (A) The binding affinity of Auranofin with SARS-CoV-2 spike was analyzed by Isothermal Titration Calorimetry. Results show binding isotherms and sensograms for the spike RBD-Auranofin interaction. (B) SDS PAGE analysis of the receptor-binding domain (RBD) of the spike protein. (C) HEK293T cells were transfected with SARS-CoV2 spike and 6 hr later 1 µM auranofin was added. Lysates were collected after 48 hr and separated by native PAGE. (D) HEK293T ACE2 cells were infected with 0.01 MOI SARS-CoV-2 in the presence of 1 µM auranofin and after 48 h infection, lysates were analyzed by Native PAGE.

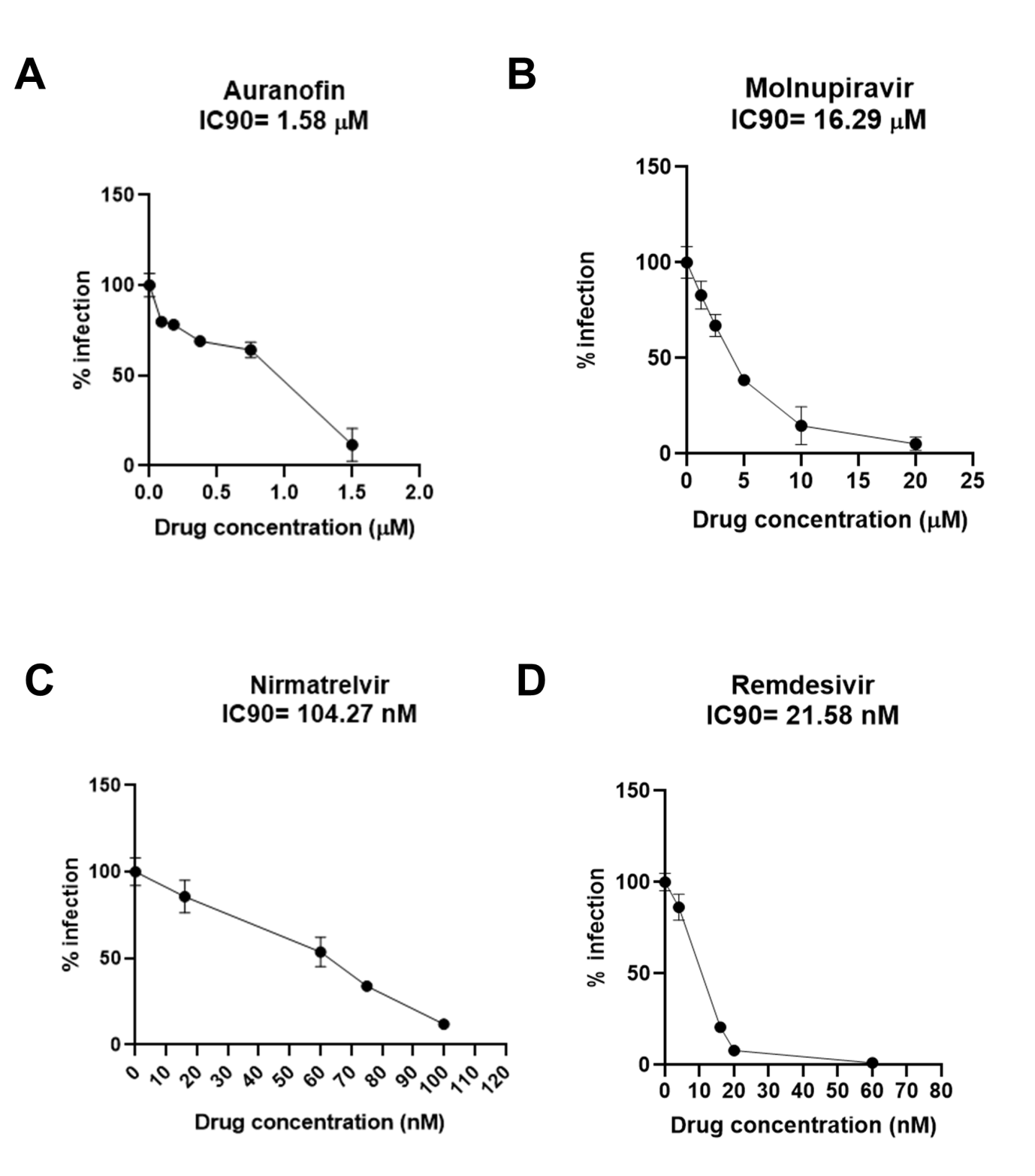

**Supplementary Fig 5. Calculation of standalone IC50 of Auranofin and DAAs.** HEK293T ACE2 cells were pre-treated with indicated concentrations of Auranofin, Molnupiravir, Nirmatrelvir, or Remdesivir for 3 h, followed by infection with 0.1 MOI SARS-CoV-2 Luc in the presence of drugs. Renilla luciferase activity was measured at 48 hpi and IC90 was calculated using Imagej/Fiji. Percentage infection compared to untreated control, showing calculated IC50 values for Auranofin, Molnupiravir, Nirmatrelvir, and Remdesivir are shown in A, B, C, and D respectively. Data represents results from 3 independent experiments. Error bars represent mean ± SD.

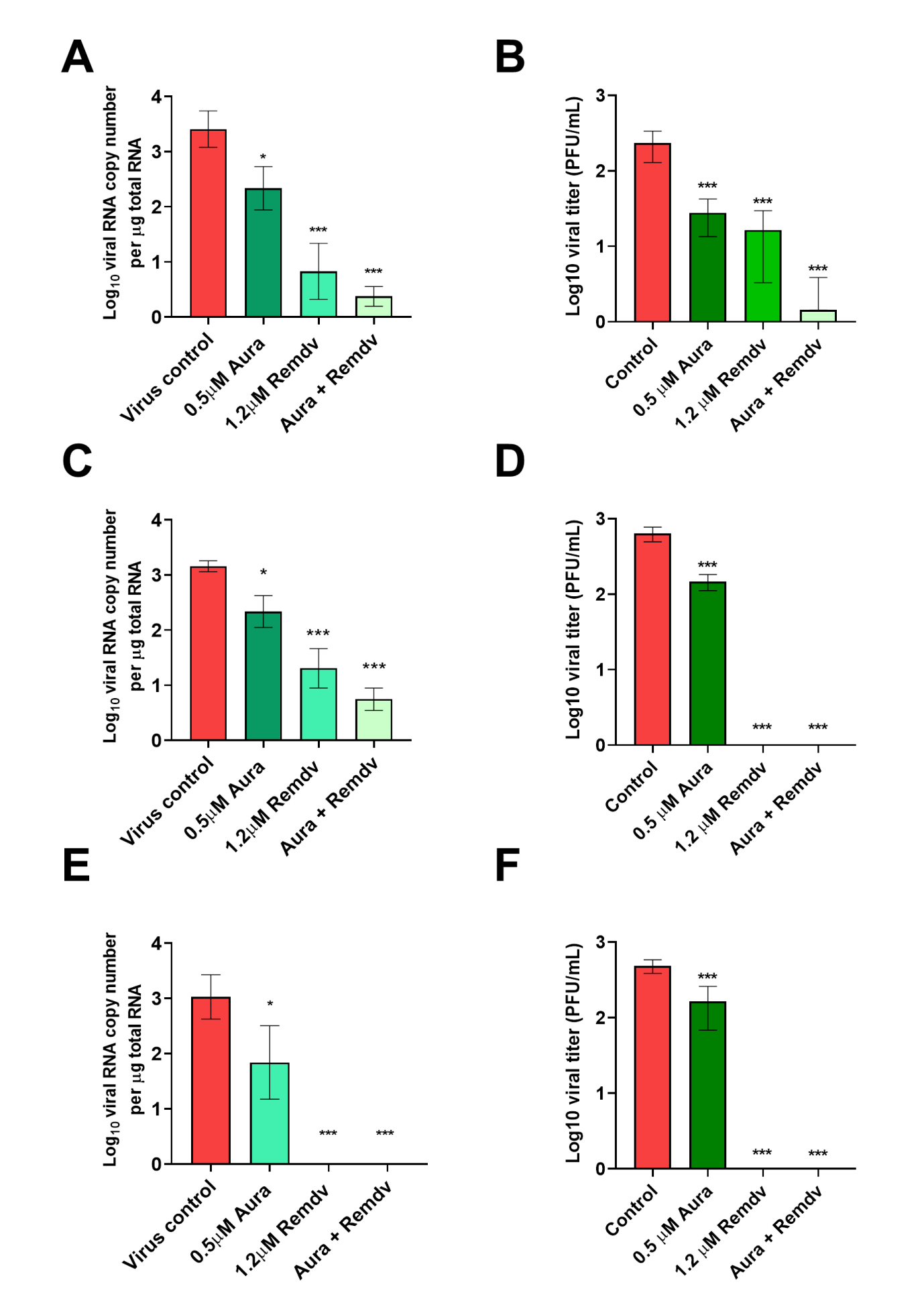

**Supplementary Figure 6. Auranofin shows synergistic antiviral activity with Remdesivir oral prodrug GS-621763.** HEK293T ACE2 cells were pretreated with Auranofin and GS-621763, either individually or in combination, as indicated. Following pretreatment, cells were infected with three different SARS-CoV-2 strains at an MOI of 0.01 in the continued presence of the respective drugs. After 48 h, cellular viral RNA levels were quantified by qRT-PCR. Infectious viral load in supernatants were quantified by plaque assay. (A,B) show quantification data for SARS-CoV-2 Hong Kong stain by qRT-PCR and plaque assay respectively. Similarly, (C,D) and (E,F) show corresponding qRT-PCR and plaque assay data for SARS-CoV-2 Delta and Omicron variants respectively. In all cases, treatment conditions were compared to the untreated virus control. All results are from 3 independent biological replicates. ∗p < 0.5, ∗∗p < 0.01, ∗∗∗p < 0.001; ns, non-significant using Welch ANOVA with Dunnett’s T3 multiple comparison tests. Error bars represent mean ± SD.
